# An AI-Cyborg System for Adaptive Intelligent Modulation of Organoid Maturation

**DOI:** 10.1101/2024.12.07.627355

**Authors:** Ren Liu, Zhaolin Ren, Xinhe Zhang, Qiang Li, Wenbo Wang, Zuwan Lin, Richard T. Lee, Jie Ding, Na Li, Jia Liu

## Abstract

Recent advancements in flexible bioelectronics have enabled continuous, long-term stable interrogation and intervention of biological systems. However, effectively utilizing the interrogated data to modulate biological systems to achieve specific biomedical and biological goals remains a challenge. In this study, we introduce an AI-driven bioelectronics system that integrates tissue-like, flexible bioelectronics with cyber learning algorithms to create a long-term, real-time bidirectional <u>b</u>ioelectronic interface with <u>o</u>ptimized <u>a</u>daptive intelligent <u>m</u>odulation (BIO-AIM). When integrated with biological systems as an AI-cyborg system, BIO-AIM continuously adapts and optimizes stimulation parameters based on stable cell state mapping, allowing for real-time, closed-loop feedback through tissue-embedded flexible electrode arrays. Applied to human pluripotent stem cell-derived cardiac organoids, BIO-AIM identifies optimized stimulation conditions that accelerate functional maturation. The effectiveness of this approach is validated through enhanced extracellular spike waveforms, increased conduction velocity, and improved sarcomere organization, outperforming both fixed and no stimulation conditions.

## Introduction

The development of bioelectronics^1-5^ that seamlessly integrate with biological systems to monitor and control tissue properties is crucial for advancing both biological studies and clinical applications. With the increasing capabilities of bioelectronics, integrating numerous sensors and actuators has become more common^6-9^. While passive recording of biological signals is well-established, interpreting these signals to generate control policies for actuators remains a challenge. The complexity of biological systems, along with limited mechanistic understanding, coupled with numerous tunable parameters for modulation, makes conventional control methods impractical.

Recent advances in artificial intelligence (AI)^10-13^ provide promising solutions. Ideally, an AI system should be given a final aim, such as achieving specific physiological states in cells and cellular networks through modulation. The AI would then learn the optimal parameters based on continuous measurements of cell properties, self-exploring to achieve the set aim.

We hypothesize that integrating bioelectronics^14-17^, AI^18-20^, and biological systems^21,22^ into an AI-cyborg system can achieve this objective. In this system, bioelectronics with sensor-actuator capabilities are controlled by AI, enabling real-time adaptive, bidirectional, and long-term stable monitoring and modulation of cellular activities across the three-dimensional (3D) tissue networks at cellular resolution.

Building such an integrated system requires overcoming three major challenges: (i) creating a bioelectronic sensing and actuating system that can seamlessly integrate in 3D with biological systems, capable of long-term stable recording and control of tissue-wide, single-cell activities without interrupting natural tissue development, differentiation, and function; (ii) developing an inference model that can continuously and accurately infer underlying cellular activities from continuous sensing data, and (iii) designing a data-driven, learning-based control system that can make decisions based on sensing data and subsequently provide optimal feedback modulations to cells through tissue-embedded actuators, precisely guiding, promoting, and ameliorating tissue-level functions and dysfunctions.

Recent advancements in bioelectronics^14-17^, computational biology^23,24^, and machine learning (ML)^18-20^, especially reinforcement-learning (RL)^25-33^, have paved the way to address these challenges. First, conventional rigid bioelectronics often cause chronic mechanical damage and inflammation when integrated with biological systems^34^. To overcome this, “tissue-like” flexible electronics have been developed using soft materials and nanoelectronics. These electronics possess properties such as tissue-level flexibility, subcellular feature size, and a mesh-like network, allowing seamless integration within 3D tissue networks^14,15^. They can record multimodal cell activities and stimulate cell activities in a chronically stable manner^15-17^, serving as a viable platform for continuous, closed-loop control.

Additionally, advances in computational methods allow for continuous inference of cell states from stable measurements. Dimensionality reduction techniques have been employed to organize cells at different states along a transition path based on dynamic genetic information, using “pseudotime”^35-37^ to trace the underlying biological progression. This approach has been previously applied to infer the functional stages of cells from their transcriptional and electrical activities^15,38^. This enables the construction of a state space for the control algorithms, where states are continuously updated with new measurements, facilitating real-time sensing and control.

Finally, RL has recently achieved much progress and has been applied to different applications across diverse domains^26,27,39-41^. Given properly defined state, action, and rewards, RL could learn an optimal feedback policy for a dynamical system through actively interacting with the system and/or a large amount of historical data^25,42^. However, implementing RL in AI-cyborg systems requires addressing two fundamental challenges: (i) providing informative states, actions, and rewards to guide RL, and (ii) overcoming RL sample efficiency challenges with limited biological experimental data^28-30^.

Here, we integrate these hardware and software innovations into an AI-driven flexible bioelectronics system, termed BIO-AIM (bioelectronics interface with optimized adaptive intelligent modulation), which provides effective modulation conditions to assist biological systems in achieving specific functional goals. As a demonstration, we applied BIO-AIM to human pluripotent stem cell-derived cardiomyocyte (hPSC-CM) organoids to promote their functional maturation. hPSC-derived organoids^43^ hold significant potential for applications such as drug screening^44^ and cell therapies^45^, but their clinical applications are often hindered by the immature state of the cells. Immature cells have a higher risk of uncontrolled proliferation, leading to potential tumor formation post-transplantation^46^ or issues with drug efficiency^47^. While there exist prior protocols^48,49^ aimed at enhancing cardiac functional maturation by continuous electrical stimulation, their effectiveness is limited since the stimulation parameters of these protocols are not adaptive to changes in the cell maturation status.

In our approach, we integrate stretchable bioelectronics with hPSC-derived organoids, forming cyborg organoids for continuous recording and stimulation through tissue-embedded electrodes. Then we leverage pseudotime, calculated from continuous recording of cardiac electrophysiological signals, for the cell state identification and reward design—by assigning higher rewards to actions leading to more advanced pseudotimes—to facilitate effective control. To address the challenge of limited sample size in biological experiments^28,29,50^, we adopt an approach based on Bayesian Optimization^51-53^, a sample-efficient black-box optimization technique. Specifically, we develop a novel parallel Bayesian Optimization methodology, building on a prior parallel Bayesian Optimization method^54^, in order to generate the control policy. Finally, control actions are delivered back to the organoids through tissue-embedded stimulation electrodes. Results show that BIO-AIM significantly enhanced maturation conditions throughout the organoid development at both cellular and tissue level (Fig. 1a), which are verified through multiple cellular and tissue level analyses.

**Fig. 1.**
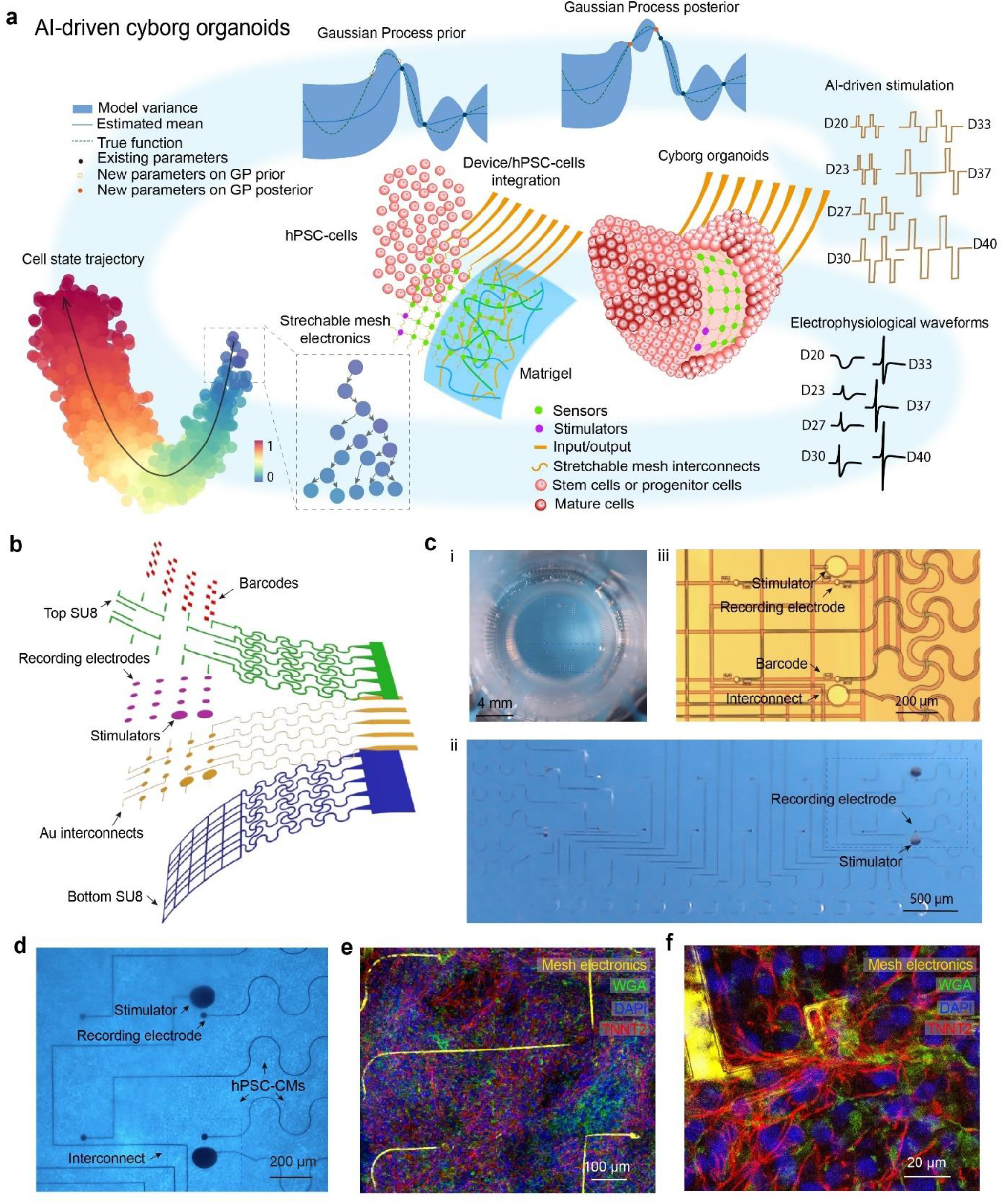
Overview of the AI-cyborg system. **a**, Schematics illustration of the AI-driven cyborg organoid workflow. Stretchable mesh electronics with sensing and actuating electrodes are embedded with human pluripotent stem cell-derived (hPSC) organoids. Electrical signals from the organoids are continuously recorded and analyzed to assess their developmental state. This state information is fed into an AI agent that utilizes Bayesian Optimization with a Gaussian Process surrogate model to learn how to accelerate the developmental transitions along the developmental trajectory by generating a control policy to stimulate organoids. The AI-generated stimulations are then applied to the organoids via the embedded actuators, iterating this closed-loop process to achieve optimized control policies that accelerate the functional maturation of hPSC-organoids. Maturation is characterized by evaluating functional and molecular phenotypes. **b**, Schematic showing the structure of stretchable mesh electronics, including polymeric passivation layers, metal interconnects, Pt electrodes, and electronic fluorescence barcodes. **c**, Representative bright-field (BF) microscopic images of stretchable mesh electronics: (i) freestanding stretchable mesh electronics with recording and stimulation electrodes in a culture chamber, (ii) zoomed-in view of the recording and stimulation electrodes highlighted in the black dashed box in (i), and (iii) representative recording and stimulation electrodes on the substrate. **d**, BF images of mesh electronics embedded in hPSC-derived cardiac organoids. **e-f**, Representative fluorescence images of immunostained 3D cyborg cardiac organoids with embedded electronics showing the sarcomere structures of hPSC-Cardiomyocytes (CMs). Red, Troponin T2 (TNNT2); green, Wheat Germ Agglutinin (WGA); blue,4′,6-diamidino-2-phenylindole (DAPI); yellow, device (SU-8 of mesh electronics were labeled by R6G).

### Creating the BIO-AIM System

We designed stretchable mesh electronics for BIO-AIM with a structure similar to previously reported designs^15,38,55,56^, including an SU-8 bottom passivation layer, 2-μm-wide gold (Au) interconnects, 25-μm-diameter platinum (Pt) sensing electrodes, and SU-8 top passivation layer, along with electronic fluorescence barcodes for sensor location registration (Fig. 1b). Previous studies^15,38,55^ have demonstrated that this design allows for stable 3D integration with organoids for long-term recording without disrupting tissue development, differentiation, and function.

To enable closed-loop control of cell activities, we incorporated individually addressable 100-μm-diameter Pt stimulation electrodes into the mesh electronics (Fig. 1b-c and Extended Data Fig. 1a-c, see Methods). Representative mesh electronics contain 64 recording electrodes and four stimulation electrodes (Fig. 1c and Extended Data Fig. 1b-c, see Methods). Pt Black was electroplated to reduce electrode impedance and increase the recording signal-to-noise ratio (SNR) (Extended Data Fig. 1d-i). The electrode demonstrated stable performance across samples (Extended Data Fig. 1d-ii) and for over 5 weeks in physiological solution (Extended Data Fig. 1d-iii), ensuring long-term electrical recording and stimulation.

We integrated stretchable mesh electronics with hPSC-CMs at 13 days of differentiation on a Matrigel layer to form 3D cyborg organoids (Fig. 1d and Extended Data Fig. 2a-b, see Methods). Immunostaining and fluorescence imaging images (Fig. 1e-f and Extended Data Fig. 2c, see Methods) confirm the tissue-wide integration of bioelectronics with cellular networks. Recording and stimulation were characterized by using a customized setup (see Methods). The stimulation controller can maintain continuous stimulation throughout the entire period of hPSC-CM development (21 days), with programmable and adaptive stimulation policies. To benchmark simultaneous stimulation and recording performance, we applied stimulation through paired stimulation electrodes within the cyborg cardiac organoids with variable amplitudes from 10 mV to 1.5 V at a frequency of 1.25 Hz. Simultaneous recording conducted across the 3D organoids under these conditions captured both stimulation spikes and cardiac electrophysiological signals, demonstrating the ability of the BIO-AIM system to simultaneously record and stimulate hPSC-CMs (Fig. 2a and Extended Data Fig. 3a-c). For stimulation amplitudes below 1.3V, the organoids maintained a ∼0.7 Hz beating frequency. At 1.5 V, the cardiac organoid signals became synchronized to the stimulation frequency, locking at 1.25 Hz. Across trials, the organoid beating frequencies remained stably synchronized at 1.25 Hz when the stimulation amplitude reached 1.5V (Extended Data Fig. 3d-e), demonstrating BIO-AIM system can modulate cardiac activity effectively.

**Fig. 2.**
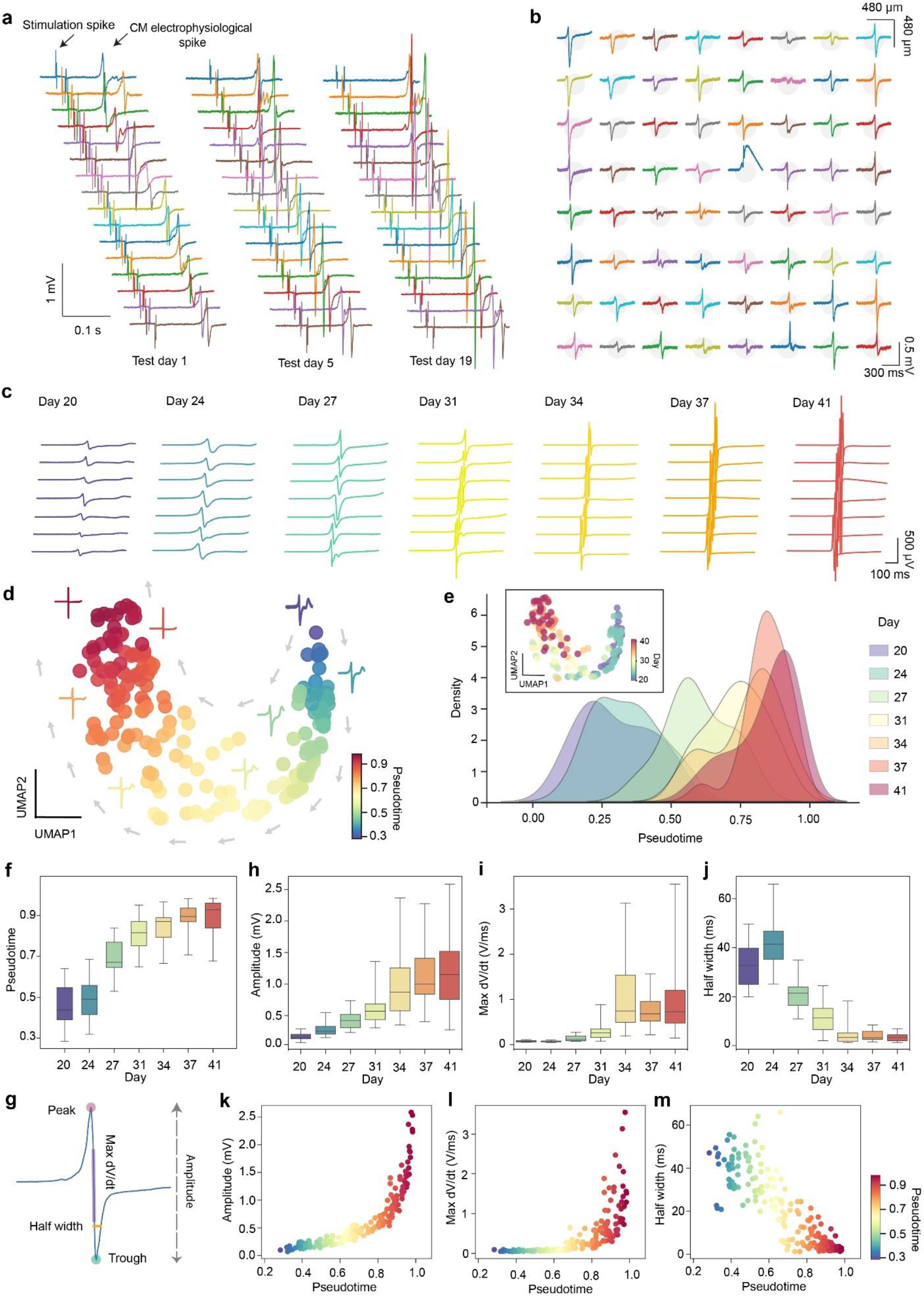
Long-term electrical recording and stimulation of cardiac activity. **a**, Representative voltage traces showing the capability of tissue-embedded sensing and stimulation electrodes for long-term tracking and stimulation of cardiac organoid electrical activity. **b**, Representative voltage traces showing electrical activities of a cardiac organoid recorded by a tissue-embedded electrode array. **c**, Representative extracellular spike waveforms recorded from a cardiac organoid at different days of cardiac differentiation. **d**, Uniform Manifold Approximation and Projection (UMAP) plot showing pseudotime-ordered electrical waveform profiles of hPSC-CMs from all channels in a representative sample at the specified differentiation days. Each dot represents an averaged electrical waveform from one channel, color-coded by inferred pseudotime. **e**, Density plot showing the distribution of pseudotime values for hPSC-CM at different differentiation days. Inset: UMAP plot showing the projection of averaged electrical waveform from each channel onto a pseudotime trajectory, highlighting the variation in cardiac activity over time. **f**, Box plots showing pseudotime values of the hPSC-CM organoids across the differentiation days. **g**, Schematic representation of the electrical features defined from a representative spike waveform for analysis. **h-j**, Box plots showing the spike waveform amplitude (**h**), max dv/dt (**i**), and half-width of the spike waveform (**j**) in hPSC-CM organoids across the differentiation days. **k-m**, Scatter plots showing the relationship between pseudotime and extracted electrical features: amplitude (**k**), maximum dv/dt (**l**), and half-width (**m**) of the spike waveforms. Each dot represents a channel-averaged electrical waveform. The data points are color-coded by pseudotime, indicating trends of increasing amplitude and maximum dV/dt, and decreasing half-width with progression in pseudotime. Box, 75% and 25% quantiles. Line, median. Whisker, 0% and 100% quantiles.

By mapping the spatial distribution of sensing electrodes across the tissue using fluorescence barcodes, we tracked cardiac electrical activity and propagation within the organoid (Fig. 2b and Extended Data Fig. 4). Electrical activity was recorded twice per week during organoid development (Fig. 2c). In a representative sample, signals were recorded on days 20, 24, 27, 31, 34, 37 and 41 of hPSC-CM differentiation, equivalent to days 7, 11, 14, 18, 21 and 24 post-reseeding onto stretchable mesh electronics (Fig. 2c). The recorded spike signals revealed clear evolution, transitioning from slow waveforms to rapid depolarization, indicative of in vitro cardiac maturation progression.

### Developing a cyber-learning framework

We developed an effective cyber-learning framework to learn optimal control policies using continuous electrical recording data to stimulate organoids through embedded stimulation electrodes, with the aim of accelerating functional maturation. The framework design involves two main steps: (i) defining the states, actions, and rewards, and (ii) designing an effective learning algorithm.

In a control framework, an agent transitions through different states over time and needs to select an action at each state. Each action generates a reward and transitions the agent to a new state in the next time step. Optimal control seeks a policy, a mapping of states to actions, that maximizes cumulative rewards over time. In our application, the agent is the organoid, the states represent the cell maturation level at each developmental stage, and the reward function is designed to accelerate maturation.

The first challenge is defining appropriate states, actions, and rewards to model the organoid system. Actions are relatively straightforward: electrical impulses delivered through embedded stimulation electrodes, represented as a 2D vector of frequency and amplitude parameters. However, determining a meaningful state from electrical recording data is challenging due to the noisy, high-dimensional nature of time-series electrical recording data.

To address this, we applied pseudotemporal trajectory inference^57^, an ML-based dimensionality reduction approach originally developed for single-cell gene expression analysis, to analyze high-dimensional electrical waveforms recorded from hPSC-CM organoids. This method reconstructs the continuous phenotypic evolution path of hPSC-CMs during functional maturation based on cell electrical spike waveforms. Previous multimodal studies pairing electrical and transcriptional measurements have shown that pseudotimporal trajectories derived from spike waveforms correspond to those from gene expression analysis, allowing for inference of cell developmental states^15^.

Using this method, we mapped cardiac electrical activities to a pseudotime value between 0 and 1, capturing the maturation level of the organoid in a compact, one-dimensional state representation. Specifically, we applied the Palantir algorithm^57^ on continuously recorded spike waveforms and used Uniform Manifold Approximation and Projection (UMAP) to project the waveforms into a 2D space, colored by inferred pseudotime values (Fig. 2d). This enables long-term electrical recording data from hPSC-CMs in organoids to be projected as an inferred pseudotemporal trajectory, mapping electrical phenotypic state transitions during cardiac development (Fig. 2e-f).

We extracted spike waveforms from each recording electrode, showing clear temporal evolution in electrical activity (Fig. 2g-j). Statistical analysis demonstrates significant increases in voltage amplitude and maximum dV/dt (Fig. 2h-i), and a significant decrease of spike half-width during hPSC-CM development (Fig. 2j). Scatter plots relating pseudotime to key electrical features reveal increasing spike amplitude (Fig. 2k), maximum dV/dt (Fig. 2l), and decreasing half-width (Fig. 2m) as pseudotime increases. These results validate the pseudotime trajectory as an accurate representation of electrical feature dynamics across cardiac developmental stages, supporting its use in explaining functional state development throughout cardiac maturation.

Finally, we defined the reward for an action at a given state (pseudotime) as the difference between the next and current pseudotime values, with the objective of achieving faster maturation. Actions that transition the organoid to a more mature state (higher pseudotime) receive higher rewards.

With states, actions and rewards specified, we designed the learning algorithm. Conventional learning-based control methods, especially RL, typically require extensive interactions with the environment and large datasets with tens of thousands to millions of data samples, often supported by high-fidelity simulators^26,58^. However, such simulators are unavailable for most biological systems^59^, including cardiac organoid development^48^. Meanwhile, real biological experiments can only yield limited samples. This constraint necessitates designing learning algorithms that can learn efficiently from limited biological data.

To address this challenge, we developed efficient, tailored learning methods that incorporate known biological principles to learn an effective stimulation policy. In a standard RL setup, it is generally assumed that the value of an action can only be evaluated at the end of an entire episode of interactions, as each action’s long-term effects must be considered^25^. Consequently, the number of samples required for an RL algorithm to learn an optimal policy scales with the length of an episode^60^. In our case, because we have limited data, we thus leverage known biological principles to design a more sample-effficient approach than generic RL. Concretely, due to the unidirectional maturation trajectory property of organoids^15,38^, we assume that actions leading to higher maturation at the subsequent timestep will also lead to improved maturation over time. This assumption allows us to evaluate actions based solely on the maturation level (pseudotime) achieved at the next transition rather than at the end of the organoid’s full maturation period. By implementing this approach, we scaled down the problem to identify the optimal stimulation action at each pseudotime, relying only on maturation level measurements at each transition. This method is more sample-efficient than standard RL approaches^25,61^.

Specifically, we adopted a Bayesian Optimization-inspired approach^51-53^ to identify effective stimulation actions at each pseudotime. We modeled changes in pseudotime between two consecutive timesteps as a function of current pseudotime and stimulation action, using a Gaussian Process model^62^. The Gaussian Process was iteratively updated with new data, and we identified the maxima of the function by selecting new points in regions with high function values (Fig. 3a). Our system allows simultaneous recording and stimulation of up to three cardiac tissues at once. Hence, to increase data efficiency, we developed a parallel Bayesian Optimization approach^54,63^ to sample new stimulation conditions for the three tissues (potentially a different one for each tissue), maximizing information gain about optimal actions to accelerate tissue maturation. The sampling strategy we use in our parallel Bayesian Optimization approach also minimizes redundancy by penalizing the selection of actions that are too similar with each other if the three tissues exhibit similar pseudotimes. To further increase data efficiency, the new data towards the Gaussian Process update may also include data from cardiac tissues with the fixed or no-stimulation protocols that are simultaneously running. Details of our learning procedure can be found in the Methods section.

**Fig. 3.**
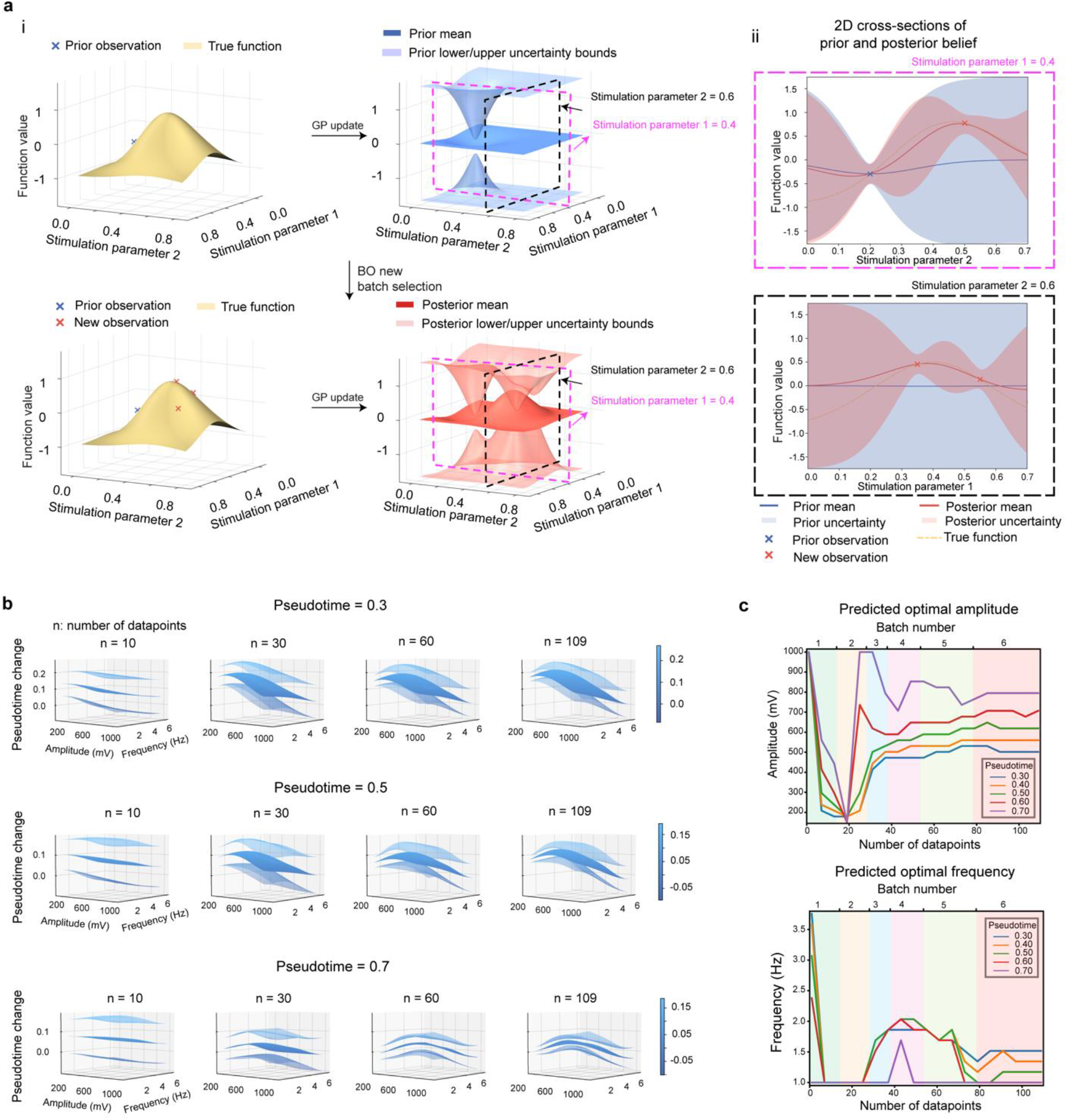
Bayesian optimization for cardiac modulation control policy generation. **a**, Schematics of the Bayesian Optimization (BO) process illustrating the update of the Gaussian Process (GP) model and selection of new batches of stimulation conditions. (i) The left subpanels show the true (simulated) function (orange surface) and (top) prior observations (blue dots) and (bottom) prior plus new observations (red dots) on the function values of an existing set of a 2D space of stimulation parameters (voltage and frequency). The right subpanels show the GP model mean (solid plane) and uncertainty bounds (shaded regions) for the function values of the two stimulation parameters before (top) and after (bottom) incorporating new observations (red dots). (ii) 2D cross-sections of the prior (blue) and posterior (red) beliefs at fixed stimulation parameters, as indicated by dashed boxes in (i). The top plot shows the cross-section at stimulation parameter 1 = 0.4, while the bottom plot shows the cross-section at stimulation parameter 2 = 0.6. Shaded regions indicate the uncertainty bounds, with the true function (dashed line) and new observations (dots) overlaid. **b**, The effect of the number of datapoints (n) and pseudotime on the prediction of function value changes (solid middle plane: GP mean function, and translucent planes: region between the two upper and lower translucent planes, GP uncertainty (± 1 SD)) across different amplitudes and frequencies. Each subplot shows the predicted change in function value as a function of amplitude (mV) and frequency (Hz) for different numbers of datapoints (n = 10, 30, 60, 100) and pseudotime values (top: 0.3, middle: 0.5, bottom: 0.7). The color scale represents the magnitude of the predicted change in function value. **c**, Predicted optimal amplitude and frequency as a function of the number of datapoints for different pseudotime values. Top and bottom plots show predicted optimal amplitude and frequency versus the number of datapoints for different pseudotime values, respectively. Lines of different colors correspond to different pseudotime values, showing the trend in optimal parameters as more data is collected.

In summary, by applying pseudotemporal trajectory analysis to continuous electrical recordings, we defined the states, actions and rewards. Using Gaussian Process and parallel Bayesian Optimization, we established a sample-efficient learning algorithm for generating cyber-control policies with limited biological data.

### Applying the BIO-AIM to organoid maturation

We integrated our cyber-learning algorithm with the cyborg organoid as an AI-cyborg organoid system. Stimulation of the organoid began one-week post-integration and continued for the entire 21-day hPSC-CM development period. Every 3-4 days (twice per week), electrical activity of the organoid was recorded to calculate the pseudotime value, which was then used to update the control policy determining the stimulation parameters through the parallel Bayesian Optimization learning strategy (see Methods).

The changes in the probabilistic Gaussian Process model (with a radial basis function (RBF) kernel) for three different fixed starting pseudotime levels are shown as a function of *n*, the number of datapoints (Fig. 3b). Each datapoint consists of an input-output pair, with the input comprising the pseudotime *s* on a measurement day and the action *a* taken starting on that day (lasting until the next measurement day), and the output being the change in pseudotime measured on the next measurement day. In each plot, the central surface indicates the expected change in pseudotime (resulting from choosing a stimulation action for a given starting pseudotime) predicted by the Gaussian Process given the current data. The translucent surfaces above and below reflect the uncertainty of the Gaussian Process in its predictions. The uncertainty in the Gaussian Process model is reduced as more data is collected for each starting pseudotime (Fig. 3b and Extended Data Fig. 5). Moreover, the mean function of the Gaussian Process gradually converges for each starting pseudotime as more data is gathered (Fig. 3b and Extended Data Fig. 5).

The Gaussian Process model’s estimated optimal stimulation parameters (frequency and amplitude) at different pseudotimes are illustrated in Figure 3c. Initially, these estimated optimal stimulation parameters fluctuate significantly due to data scarcity; however, as more data is collected, they begin to converge to a distinct level for each pseudotime (Fig. 3c). Notably, the model suggests that higher stimulation amplitudes are optimal at higher pseudotimes, while lower stimulation frequencies are optimal at lower pseudotimes.

To evaluate the effectiveness of BIO-AIM in accelerating organoid maturation, the cyborg organoids were divided into three groups (Fig. 4a): a control group integrated with flexible electronics for chronic recording but without stimulation, a group with fixed stimulation parameters from previous studies optimized for cardiac organoid maturation^48,49^, and a group with an AI-generated stimulation parameters that dynamically adjusted in response to real-time cardiac electrical signals as described above. Notably, the BIO-AIM-generated stimulation parameters resulted in a unique pattern, distinct from the fixed and no-stimulation conditions (Fig. 4b), suggesting that varied stimulation amplitudes and frequencies differentially affect cardiac tissue electrical properties. Importantly, the AI-driven group demonstrates more mature electrical activity patterns, as shown by representative waveforms throughout development (Fig. 4c).

**Fig. 4.**
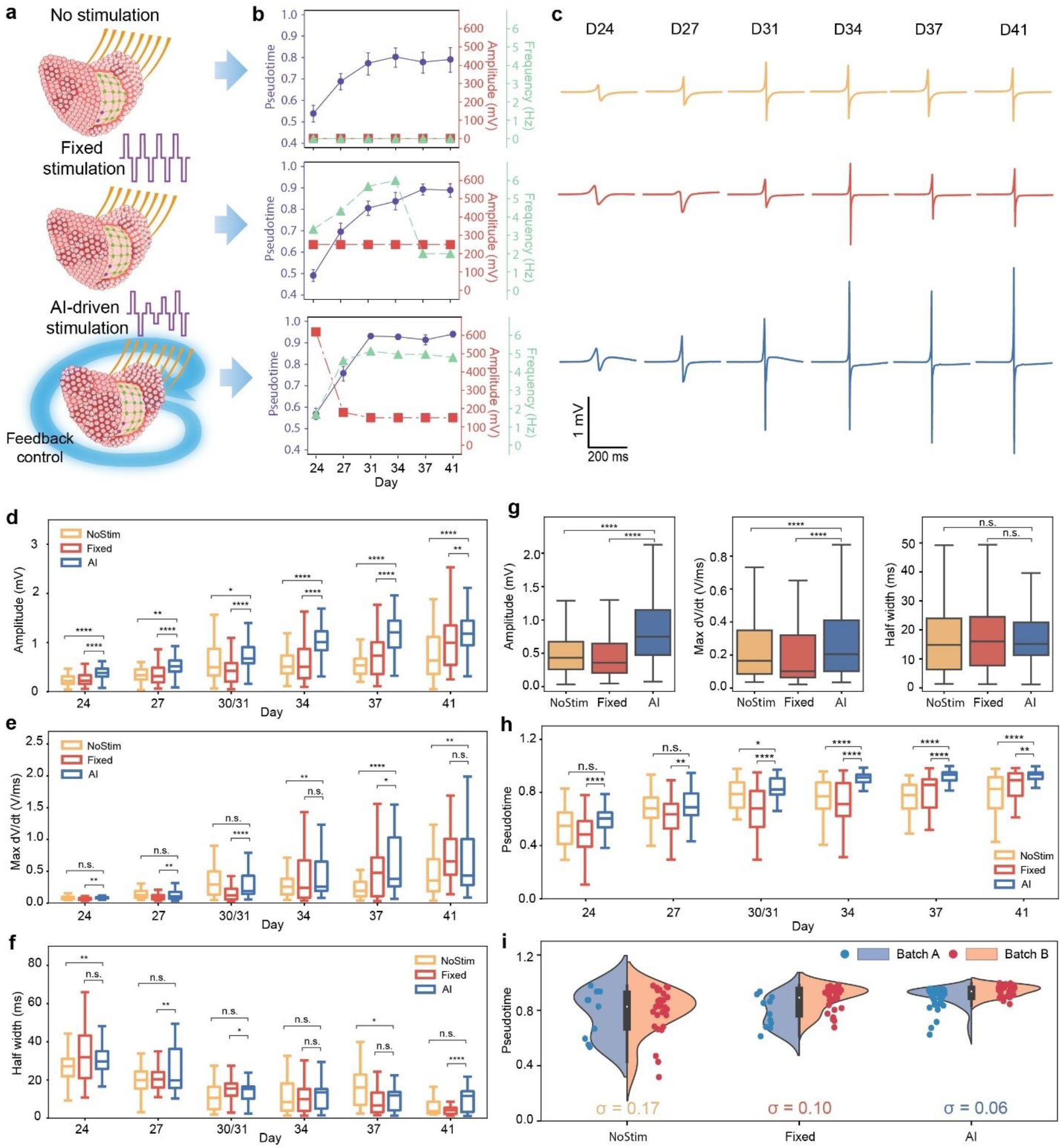
Advancing hPSC-CM functional maturation through the AI-cyborg system. **a**, Schematics of three stimulation strategies: no stimulation (NoStim), fixed stimulation (Fixed), and AI-driven stimulation (AI). **b**, Line graph showing the statistical changes in pseudotime values, as well as the amplitudes and frequencies applied during stimulation across different differentiation days. Values are means ± 95% confidence intervals. **c**, Representative raw voltage spike waveforms recorded at different differentiation days for no, fixed, and AI-driven stimulation groups. **d-f**, Box plots comparing the spike waveform features for no, fixed and AI-drive stimulation groups across different days of differentiation: **d**, amplitudes, **e**, maximum dV/dt values and **f**, half widths. **** p < 0.0001, ** p < 0.01, * p < 0.05, n.s. not significant. Box, 75% and 25% quantiles. Line, median. Whisker, the most extreme data point within the median ± 1.5 interquartile range (IQR). **g**, Box plots comparing the electrical properties of cardiac organoids under different stimulation policies: (Left) Amplitude of the spike waveform. (Middle) Maximum dV/dt of the spike waveform. (Right) Half width of the spike waveform. **h**, Box plots showing the pseudotime values for no stimulation, fixed stimulation, and AI-driven stimulation groups across different differentiation days. **** p<0.0001, ** p<0.01, * p<0.05, n.s. not significant. Box, 75% and 25% quantiles. Line, median. Whisker, the most extreme data point within the median ± 1.5 IQR. **i**, Violin plots comparing the pseudotime distribution on 41 days of differentiation from two batches of samples for three stimulation groups. SDs (σ) are computed across the two batches.

We further analyzed key electrical features of the extracellular field potential waveform, including amplitude, maximum dV/dt, and half-width across all groups (Fig. 4d-f). The AI-driven group consistently exhibited statistically significant increases in voltage amplitude and maximum dV/dt compared to fixed and no-stimulation group on each recording day (Fig. 4d-e), with occasional increases in spike half-width (Fig. 4f). Overall, the AI-driven group achieved statistically significantly higher amplitude and maximum dV/dt while maintaining a similar half-width (Fig. 4g). Previous studies have confirmed that these features correlate with molecular phenotypes and electrical signal propagation, serving as biomarkers of electrophysiological maturity for assessing the functional state of cardiac organoids^15,38,64^, suggesting a greater maturity AI-driven group electrical phenotype.

Pseudotime progression over time illustrates that AI-driven stimulation significantly accelerates maturation compared to the fixed and no-stimulation groups (Fig. 4h). Cardiac electrical activities across groups were visualized using UMAP (Extended Data Fig. 6a), with the AI-driven group displaying an extended pseudotime trajectory (Extended Data Fig. 6b). The density plots show that the AI-driven approach results in a more advanced progression along the pseudotime trajectory compared to fixed and no-stimulation groups (Extended Data Fig. 6c). We next investigated the correlation between pseudotime with key electrical features of of the extracellular field potential waveform, including amplitude, maximum dV/dt, and half-width across the three groups. We projected the electrical features from three different groups of cells on the joint UMAP embedding, which demonstrates the effects of AI-driven stimulation enhance cardiac organoid maturation process across the developmental trajectory (Extended Data Fig. 6d).

We analyzed the pseudotime distribution in later developmental stages across different batches (Fig. 4i). The AI-driven group demonstrated statistically significant reduction in variance across batches, indicating more uniform electrical function across the tissue compared to fixed and no-stimulation groups. This validates that the AI-driven stimulation can further reduce the batch-to-batch variation in cardiac organoid maturation.

Beyond accelerated cellular electrical maturation, we investigated whether AI-driven stimulation would enhance network functions at the tissue level. Using the tissue-embedded recording electrode array, we chronically mapped 2D/3D electrical signals across the organoids, which enabled us to calculate electrical signal propagation using activation mapping (Fig. 5 and Extended Data Fig. 7-8). Voltage traces of spikes from electrodes distributed across tissues show a gradual reduction in spike latency over time in all three groups (Fig. 5a, Extended Data Fig. 7a, and Extended Data Fig. 8a), with the AI-driven group exhibiting the most significant decrease in temporal delays. Heatmaps of propagation delay (Fig. 5b, Extended Data Fig. 7b, and Extended Data Fig. 8b) confirm that AI-driven group consistently achieved faster propagation. Statistical analysis of conduction velocity (see Methods)^65,66^ further demonstrates a significantly higher increase in the AI-driven group compared to fixed and no-stimulation groups (Fig. 5c, Extended Data Fig. 7c, and Extended Data Fig. 8c). Specifically, AI-driven stimulation group reached a mean velocity of 102.6 ± 2.3 mm/s (mean ± SEM), compared to 60.6 **±** 1.5 mm/s for the fixed stimulation group (Fig. 5c). These results demonstrate that BIO-AIM-enabled AI-driven stimulation significantly enhances synchronization of cell network activity and accelerates signal propagation, promoting tissue-level functional maturation.

**Fig. 5.**
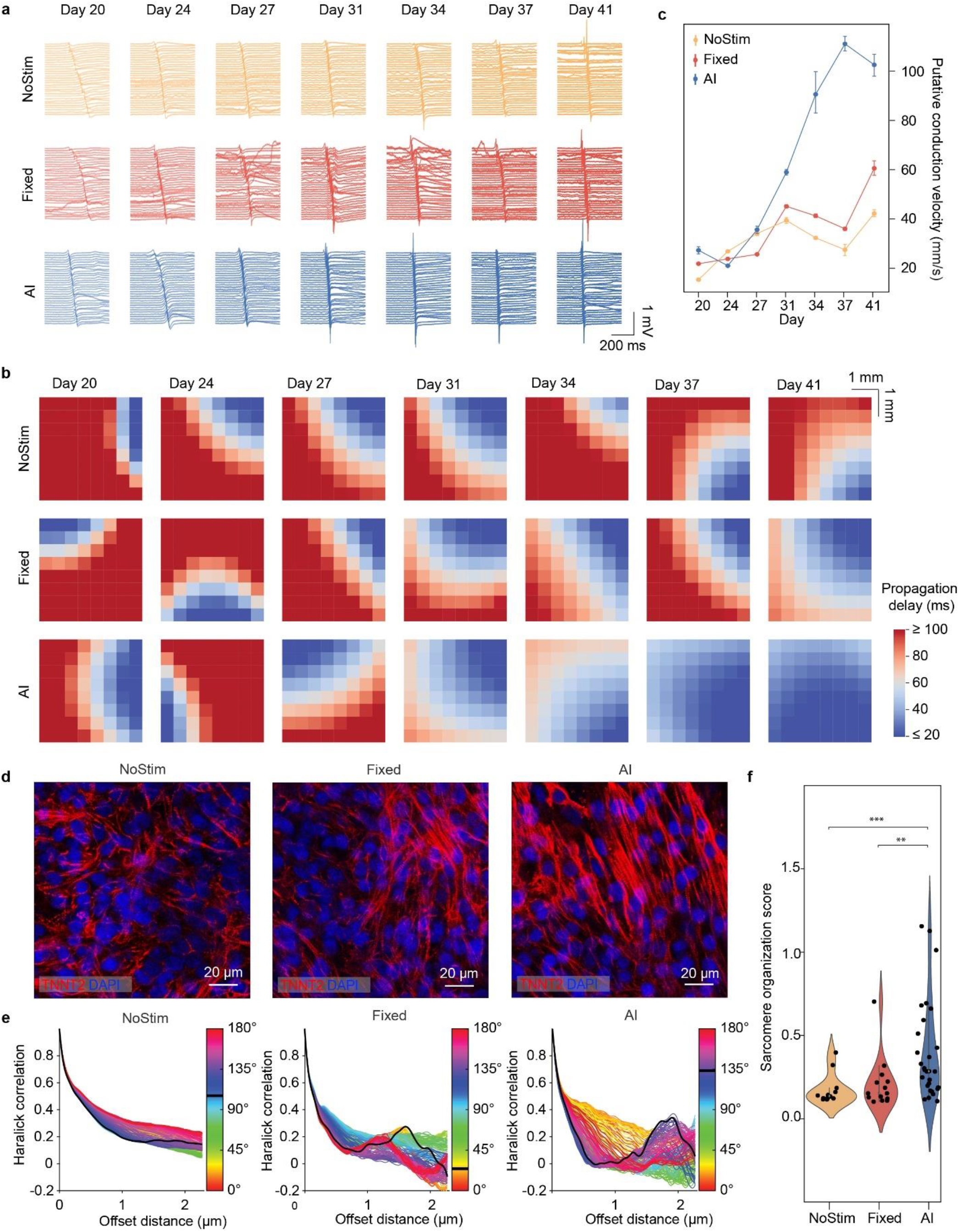
Functional maturation characterizations in cardiac organoids. **a**, Representative raw voltage traces showing electrical activity mapping of cardiac organoids under different stimulation conditions across differentiation days. **b**, Heatmaps illustrating the putative propagation delay of electrical signals from the electrical mapping, projected onto the recording electrode arrangement for the three stimulation groups from the same batch of cells at the same differentiation days. **c**, Line graph showing the statistical changes in putative conduction velocity over channels for three stimulation groups across different differentiation days. Values are mean values ± 95% confidence intervals. **d**, Confocal fluorescence images of immunostained cardiac organoids at 65 days of differentiation from three conditions: Red, TNNT2 and blue, DAPI. **e**, Plots showing the haralick correlation as a function of offset distance for representative cells of the three stimulation groups. Colors represent different angles, with the color scale indicating the degree of correlation. **f**, Violin plots comparing the sarcomere organization score for three stimulation groups. *** p<0.001, ** p<0.01, unpaired two tailed t-test.

Additionally, we evaluated sarcomere alignment, which is crucial for effective muscle contraction, a key indicator of cardiac tissue maturation^48^. Immunostaining for cardiac troponin T (TNNT2) showed poorly organized sarcomere structures in no-stimulation group, moderate organization in the fixed stimulation group, and well-aligned, organized sarcomere structures in the AI-driven group (Fig. 5d and Extended Data Fig. 9). The analysis of sarcomere organization using Haralick correlation^67^ confirmed that AI-driven stimulation statistically significantly improves sarcomere organization score (Fig. 5e-f). Together, these results collectively demonstrate the effectiveness of BIO-AIM in enhancing cardiac structural maturation.

## Discussion

In this work, we integrated tissue-like bioelectronics with cyber-learning algorithms to create BIO-AIM, an AI-driven, long-term bidirectional bioelectronic interface capable of continuously adapting and optimizing control policies based on stable cell state mapping. When applied to cardiac organoids, BIO-AIM allows AI-driven closed-loop feedback through tissue-embedded flexible electrode arrays, identifying effective stimulation conditions to accelerate organoid functional maturation. This accelerated maturation is validated by enhanced extracellular spike waveform features, increased conduction velocity, and improved sarcomere organization compared to fixed and no-stimulation controls.

From a biological perspective, natural organ development involves nerve innervation that provides closed-loop sensing and stimulation throughout in vivo development—an aspect typically lacking in in vitro tissue systems. BIO-AIM mimics nerve innervation by continuously monitoring and controlling the organoid system, promoting more accurate cell differentiation, tissue organization, and overall development. This system can be applied to various organoid types and tissue models, potentially advancing our understanding of complex diseases, developing new treatments, and even enabling applications in vivo, which could significantly impact tissue engineering and regenerative medicine. In addition, the adaptive learning capabilities of the AI system ensure continuous improvement over time, enhancing its performance and effectiveness.

During our experiments, the cyber-learning algorithm received only tissue functional states as input and autonomously explores the stimulation parameters. After multiple rounds of exploration, the algorithm identifies optimized, dynamically adaptive stimulation parameters. Further studies are needed to explain the AI-learned stimulation policy, understand why it accelerates functional maturation, and determine the extent to which this system can enhance tissue development. Detailed analysis of gene and protein expression, as well as characterizations of cell connectivity and morphological, may be useful in understanding the mechanisms behind the AI-driven maturation acceleration.

Further development in both hardware and software is needed to enhance the performance of this system. Hardware improvements could involve scaling up the number and density of sensors and actuators through the integration of advanced fabrication techniques and application-specific integrated circuits (ASIC) systems^8,9^ to increase the resolution and accuracy of the system.

Software enhancements could focus on the following aspects of the RL component. Firstly, currently, we use a Gaussian Process to model the dynamics of the cardiac organoid, which is suitable for low-data regimes^62^. However, as more data becomes available, representation-learning techniques^68-71^ could be employed to model biological dynamics more accurately.

Secondly, the current system views an organoid as a single entity, suggesting a single stimulation across all channels. In more complex tissues, such as brain organoids^14,56^, where different cell types coexist, a network policy that provides different stimulation parameters to different channels and locations may be necessary. Coordinating such a network policy will require the development of multi-agent networked RL techniques^54^. Thirdly, for more complex tissues, long-term effects of actions may need to be considered, potentially requiring adaptation of techniques to handle the challenge of higher sample complexity^60^. Finally, multifunctional sensors and stimulators could be incorporated to provide multimodal input (e.g., cell molecular or structure information) and multimodal stimulation (e.g., chemical and mechanical) to the tissue^3,5,72^.

Developing a multimodal AI agent capable of efficiently integrating different modalities^15,17^ will be essential to achieving this.

## Methods

### Device fabrication and packaging

#### 1. Stretchable mesh electronics fabrication

The ultra-flexible, stretchable mesh nanoelectronics were fabricated following established methods^38,73,74^. Clean 4-inch double-side polished glass wafers were used as substrates, prepared using piranha solution, rinsed with deionized (DI) water, and dried with nitrogen. Hexamethyldisilazane (HMDS, MicroChem) was spin-coated at 4000 rpm. A bilayer of LOR 3A (300 nm, MicroChem) and S1805 (500 nm, MicroChem) was then spin-coated at 4000 rpm, followed by baking at 180 °C for 5 min and 115 °C for 1 min, respectively. Ni patterns were exposed by Ultraviolet (UV) light (Karl Suss MA6 mask aligner) at 40 mJ/cm^2^ and developed using CD-26 developer (MICROPOSIT) to define nickel (Ni) patterns. O_2_ plasma treatment (Anatech Barrel Plasma System) was applied to remove residues. A 100-nm-thick Ni layer was deposited using a Sharon Thermal Evaporator, followed by a standard lift-off procedure in remover PG (MicroChem). Next, SU-8 2000.5 precursor (MicroChem) was spin-coated at 4000 rpm, pre-baked at 65 °C/95 °C for 2 min each, exposed to UV light at 200 mJ/cm^2^, post-baked at 65 °C/95 °C for 2 min each, developed using SU-8 developer (MicroChem), rinsed by isopropyl alcohol (IPA), dried with nitrogen, and hard-baked at 180 °C for 40 min. Then, patterns of interconnects were defined using HMDS/LOR3A/S1805 bilayer photoresists as described above, followed by depositing 5/40/5-nm-thick chromium/gold/chromium (Cr/Au/Cr) using an electron-beam evaporator (Denton). A standard lift-off procedure in remover PG (MicroChem) was used again. Patterns of electrode arrays were defined using the same method for fabricating the interconnects as described above. 5/50-nm-thick chromium/platinum (Cr/Pt) was deposited using the electron-beam evaporator, followed by the same lift-off procedure. Finally, a top SU-8 encapsulating layer was defined using the same method for the bottom SU-8. A layer of fluorescence barcodes was patterned with 0.004 wt‰ Rhodamin 6G powder (Sigma-Aldrich) added into SU-8 precursor.

#### 2. Stretchable mesh electronics packaging

The flexible flat cable (Molex) was soldered onto the input/output (I/O) pads using a flip-chip bonder (Finetech Fineplacer). A culture chamber was then glued to the substrate, fully enclosing the device with a biocompatible adhesive (Kwik-Sil, WPI). Then, Pt black (PtB) was electroplated on the Pt electrodes. The device was rinsed with deionized (DI) water for 30 s and dried by nitrogen. Finally, the surface of the device was treated with light oxygen plasma (Anatech 106 oxygen plasma barrel asher), followed by adding 1 mL of Ni etchant (type TFB, Transene) into the chamber for 2 to 4 hours to completely release the device from the substrate. The device was then prepared for sterilization before cell culture.

### Tissue experiments

#### 1. Differentiation of human pluripotent stem cells (hPSCs)-derived cardiomyocytes

HPSCs (hPSC-line GSB-L88, Greenstone) were seeded into tissue culture plates and allowed to grow for 4 days. Cardiomyocytes (CMs) were then differentiated from hPSCs following previous protocols^15,55,75,76^. On day 0, the media was changed to RPMI 1640 + B27-insulin (RPMI/B27-insulin) containing the GSK3 inhibitor CHIR99021 (6 µM, Selleck Chemicals). On day 1, the media was changed to RPMI 1640 /B27-insulin. On day 3, the media was changed to RPMI 1640/B27-insulin containing IWR1 (5 µM). On day 5, the media was changed to fresh RPMI/B27-insulin. On day 7, the media was changed to RPMI 1640/B27 (10 µg/mL), with media subsequently changed every 2-3 days with RPMI 1640/B27. Beating CMs were typically observed between days 7-9 of differentiation.

#### 2. 3D cyborg cardiac organoid culture

The released stretchable mesh electronics were first rinsed with DI water and decontaminated by 70% ethanol. Then the device was incubated with Poly-D-lysine hydrobromide (0.01% w/v) overnight and subsequently coated with Matrigel solution (10 mg/mL) for 1 hour at 37 °C. The hPSC-derived CMs (hPSC-CMs) were seeded onto the stretchable mesh nanoelectronics placed on the surface of the Matrigel layer to form 3D cyborg cardiac organoids. Firstly, the device was pre-cooled on ice, and 60 µL Matrigel solution (10 mg/mL) was added to cover the whole bottom of the cell culture chamber. The device was incubated at 37 °C to solidify the Matrigel. Secondly, the 2D cultured hPSC-CMs were dissociated into single cells. About 3∼4 million cells were suspended in 1 mL RPMI 1640 medium plus 1% B27 and transferred onto the cured Matrigel/electronics substrate, maintaining at 37 °C, 5% CO_2_. A 5 μM rock inhibitor (Y27632) was added on the first day to improve cell viability. Finally, the 3D cyborg cardiac organoid was maintained at 37 °C, 5% CO_2_ in the incubator, with medium changed daily.

#### 3. Immunostaining and imaging

Immunostaining was performed as previously reported^15,38,55^ Briefly, samples were placed in an X-CLARITY hydrogel polymerization device for 4 hours at 37 °C with -90 kPa vacuum and then placed in the X-CLARITY electrophoretic tissue clearing (ETC) chamber to extract electrophoretic lipids. For staining, primary antibodies TNNT2 and WAG were incubated at 4 °C for 4 days, followed by secondary antibodies at 4°C for 2 days. Samples were submerged in an optical clearing solution and embedded in 1% agarose gel before imaging using the Leica TCS SP8 confocal microscope.

### Electronics characterization, recording, and stimulation

#### 1. Device characterization

The electrochemical impedance spectroscopy (EIS) of the electrodes in stretchable mesh electronics was conducted following previous methods^15,55^. A three-electrode setup was used to measure the EIS with a standard silver/silver chloride (Ag/AgCl) electrode as the reference and a platinum wire as the counter electrode. The device was immersed in phosphate-buffered saline (PBS) solution during measurement. EIS data were collected using SP-150 potentiostat (BioLogic) with EC-lab software.

#### 2. Recording and stimulation

Electrical activity was recorded using an RHD 64-channel headstage (Intan technologies) connected to the Intan RHD recording controller (Intan technologies). A custom-made printed circuit board with an integrated Omnetics connector (Omnetics) linked the RHD 64-channel headstage to the device via an FFC connector (Digi-Key). The organoid culture media was grounded, with a reference electrode far from the device and cells. Platinum wires were used for both ground and reference electrodes. During measurement, cyborg cardiac organoids were placed on a battery-powered warming plate, maintaining thermostatic 37 °C throughout the recoding. Electrophysiological recordings were captured at a 20 kHz sampling rate. Electrophysiological activities were recorded twice a week. For stimulation, samples were maintained under the electrical stimulation using a Master-8 (AMPI), with wire connected to the samples inside the incubator.

### Data analysis

#### 1. Electrical data analysis

We used a customized Python (version 3.8) pipeline for electrical waveform extract and downstream feature analysis. Raw recording traces were first smoothed using the “gaussian_filter” function, and spike candidates were identified using the “find_peaks” function from the Scipy library. A threshold of 5 times the standard deviation (SD) from the mean of the smoothed traces was applied during peak detection. Spike candidates from the same recording channels were averaged to obtain spike templates, which were manually curated to remove noises. These spike templates were used to compute waveform features: peak (maximum recorded voltage), trough (minimum recorded voltage), and amplitude (voltage difference between peak and trough). The maximum dV/dt was calculated as the maximum absolute derivative value of the spike templates. The half-width was calculated as the time difference between the time points where half of the trough value was first and last reached.

We inferred pseudotime^77,78^ scores from the spike templates using the Palantir algorithm within the Scanpy package. Spike templates from all samples across all experiment days were combined as the input to the “palantir” function to infer the pseudotime scores. The UMAP was used to project all the spike templates onto a 2D space for visualizing the pseudotime distribution.

#### 2. Sarcomere alignment quantification

The Sarcomere Organization Texture Analysis algorithm was applied, using Haralick texture structures to quantify sarcomere alignment as previously introduced^38,67^.

### 3. Conduction velocity computation

Conduction velocity was computed based on previous methods with some modifications^65,66^. Firstly, a polynomial surface was fitted to the space-time coordinates of recorded electrical activities. The fitted time lag and distance of each channel with respect to the earliest beat identified were calculated and averaged to obtain the conduction velocity for each detected event.

### Cyber-Learning Model

#### 1. Learning algorithm summary

We hypothesized that appropriately stimulating the cardiac tissue with electrical impulses can accelerate its maturation process based on previous reports^48,49^. A Bayesian Optimization (BO)^51^ based Reinforcement Learning (RL) approach was used to find an optimal electrical stimulation policy that provide an adaptive stimulation depending on tissue maturation level.

#### 2. Problem setup and preliminaries

Let 𝒮: = [0,1] denote the state-space representing the pseudotime. For the action space, the following constraints are imposed on any valid physical action *ā*: any *ā* = (*ā*_1_, *ā*_2_) should satisfy (i) *ā*_1_Hz lies between 1Hz and 6Hz, (ii) *ā*_2_ mV lies between 150mV to 1000mV. These constraints have been imposed due to our prior knowledge of the physical dynamics. Any valid action *ā* is mapped to an action *a* in a two-dimensional action space 𝒜 = [0,1] × [0,1] which is the product of two unit intervals, via the map

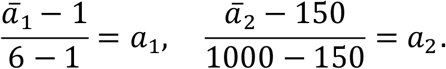

Thus, the action space is represented as 𝒜 = [0,1] × [0,1].

Let *s* denote the pseudotime of a cardiac device measured at a day *t*. Let *a* be the action applied to the cardiac device starting on day *t*. Let *t*′ > *t* be the next day where a measurement is taken; the action *a* is continuously applied to the device until day *t*′. In our experiments, the gap between *t*′ and *t* is either 3 or 4. Let *s*′ be the pseudotime of the same cardiac device when measured on day *t*′. We assume that there exists some map *f*: 𝒮 × 𝒜 → ℝ such that

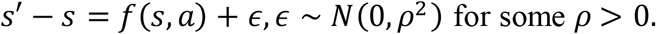

Hence, the change in pseudotime between day *t* and day *t*′ (day of next measurement) can be explained (up to some noise term ϵ) by a function *f*(*s, a*) that takes into account the pseudotime *s* on day *t*, as well as the action *a* taken between day *t* and day *t*′. Then, for any pseudotime *s*, we aimed to learn an action *a*^*^(*s*) ∈ 𝒜 such that *a*^*^(*s*) ∈ arg max_*a*_*f*(*s, a*). Then, *a*^*^(*s*) will be the action that advances the next pseudotime *s*′ the most (in expectation).

To learn *a*^*^(*s*) across the different *s* ∈ [0,1], the function *f*(*s, a*) is modelled by a Gaussian Process (GP), and then the function is optimized across varying *s* via Bayesian Optimization (BO). We assume that *f*(*s, a*) is drawn from a GP prior, i.e. *f*(*z*) ∼ *GP*(0, *k*(*z, z*′)) where *z* : = (*s, a*) and *z*′ : = (*s*′, *a*′) for another state-action pair (*s*′, *a*′), and *k*(*z, z*′) denotes an initial kernel. By using a GP, the posterior belief about *f*(*s, a*) is adaptively updated each time new data is collected.

In each experiment, let *m* denote the batchsize, which is the total number of devices being maintained in the experiment. On each measurement day *t*, the pseudotimes 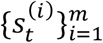 for each of the *m* devices are measured, and *m* different actions 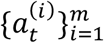 are picked. The next pseudotimes 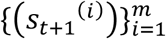 are then measured at the next measurement day.

We refer to any state-action tuple *z* = (*s, a*) as a point in the set 𝒵 := 𝒮 × 𝒜, and refer to the actual measured *s*′ − *s* at the next measurement as an evaluation of the point *z* = (*s, a*). Let 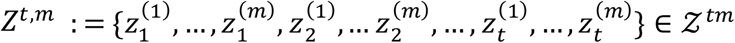 denote the *tm* points available after *t* rounds where *m* points (1 point per device) were evaluated per round, with 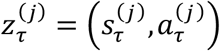 denoting the *j*-th point evaluated at the τ-th round; in the rest of the paper, we omit the dependence on the batchsize *m* and refer to *Z*^*t*,*m*^ as *Z*^*t*^. Let **y**_*t*_ denotes 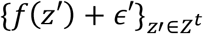, where ϵ′ ∼ *N*(0, *ρ*^2^), and ℱ_*t*_ : = {*Z*^*t*^, **y**_*t*_}. Given data ℱ_*t*_, for any *z* ∈ 𝒵 : = 𝒮 × 𝒜, the GP can be described as *f* ∣ ℱ_*t*_ ∼ *GP*(*μ*_*t*_(*z*), *k*_*t*_(*z, z*′)), where

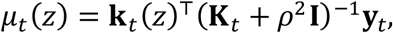

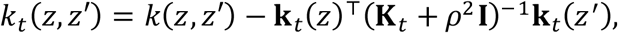

with 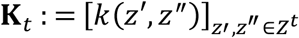 denoting the empirical kernel matrix, 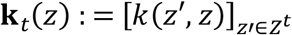 . For any *z* ∈ 𝒵, the GP belief at *z* satisfies 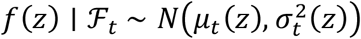, where the posterior variance is

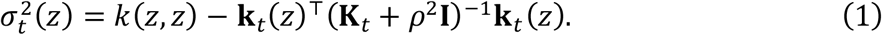

Given any set of points and corresponding measurements ℱ : = {*Z*, **y**}, if

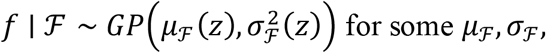

we denote *μ*_ℱ_ (*z*) as *μ*(*z*) ∣ ℱ and *σ*_ℱ_(*z*) as *σ*(*z*) ∣ ℱ.

For any set of *B* points {*z*^(*b*)^}_*b*∈[*B*]_ ∈ 𝒵, we denote

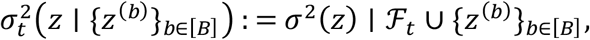

which can equivalently be expressed as

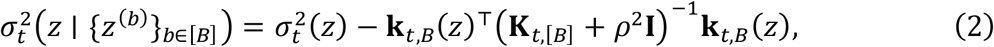

where 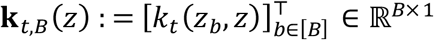, and 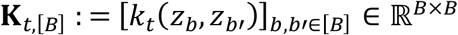.

Next, we describe the algorithm that we use to learn *a*^*^(*s*) across varying *s* ∈ 𝒮 = [0,1] in detail.

#### 3. Algorithm

Our algorithm is described in Algorithm 1 below. Lines 1 through 4 comprise the initialization of the algorithm. The initial kernel is picked to be the Matérn kernel^62^ with *ν* = 1.5, the noise variance estimate is *ρ*^2^ = ln 2, and the confidence parameter sequence is *β*_*t*_ ≡ 0.2.

Next, Lines 5 through 13 describe the GP update, action selection, and data update in each round. In Line 6, the posterior mean and variance of the GP is updated given the current data. Next, in Line 7, the pseudotimes of the active devices in the round are measured. The sampling strategy is described in Lines 8 through 10. The details of the sampling strategy for the action selection in Algorithm 1 are as follows.

##### Algorithm 1

Bayesian Learning algorithm for learning cyber-layer policy

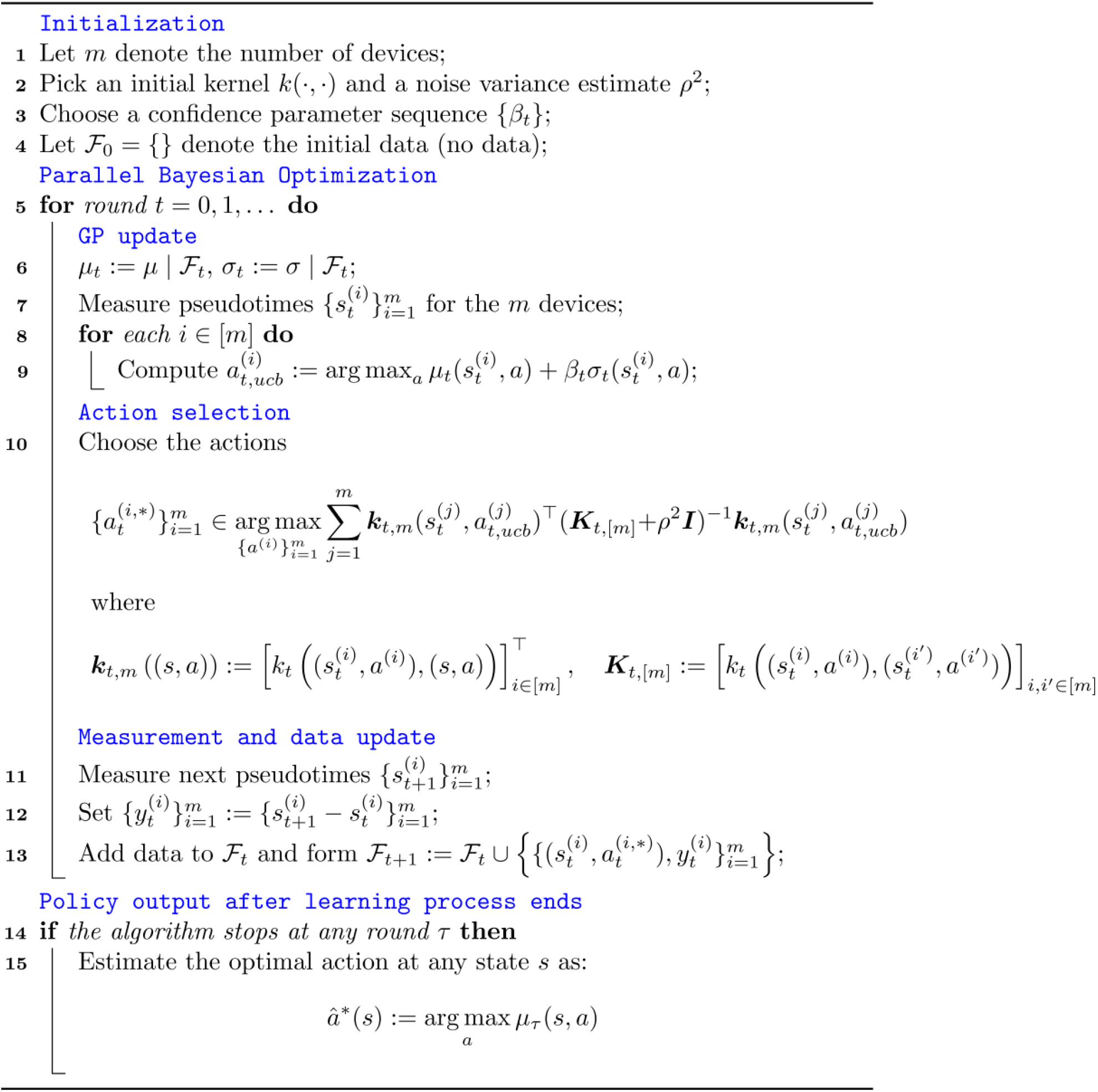

In the action sampling strategy, the max-value Entropy Search (ES)^52^ framework is adapted to the parallel BO setting. In this setting, the max-value ES sampling strategy takes the form

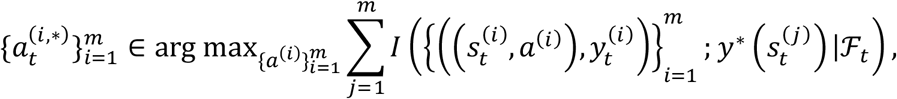

where 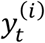 denotes the measured outcome 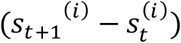 (which is a random variable), 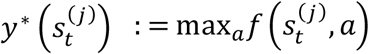 denotes the maximal value of *f* holding 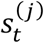 fixed, and for any random variables *X, Y* and sigma-algebra ℱ, *I*(*X; Y* ∣ ℱ) denotes the mutual information between *X* and *Y* given ℱ. It has previously^53^ been shown that such a sampling strategy is an efficient approximation for choosing a collection of actions 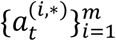 such that, given the existing information ℱ_*t*_, the potential information gain about the optimal actions at the states in the current round 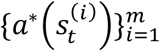, which are random variables given ℱ_*t*_, is maximized. However, the sampling function for max-value ES is computationally expensive to optimize in the parallel BO setting^54^. Thus, instead the following approximation to max-value ES for the parallel BO setting is used^54^:

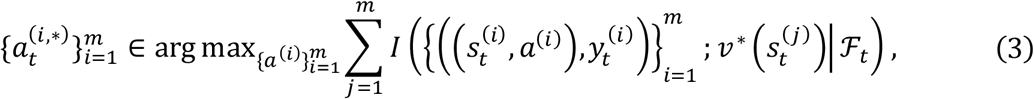

where

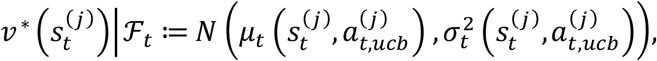

and 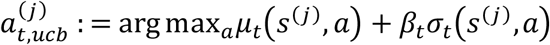 with *β*_*t*_ denoting a confidence parameter. The action 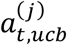 is an action that maximizes an upper-confidence-bound (UCB), holding 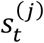 fixed, of the function 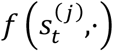 given the current knowledge ℱ_*t*_ . In the objective above, the 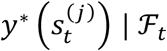 term, which appears in max-value ES, is approximated by an estimate 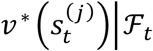 which is the pointwise Gaussian 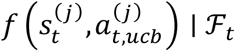.

In addition, define

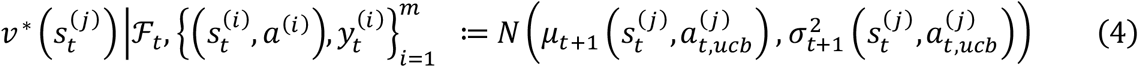

where 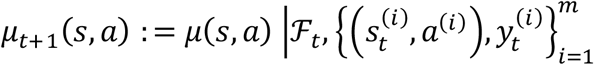, and

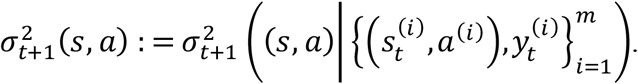

The objective in (3) can then be further simplified as follows:

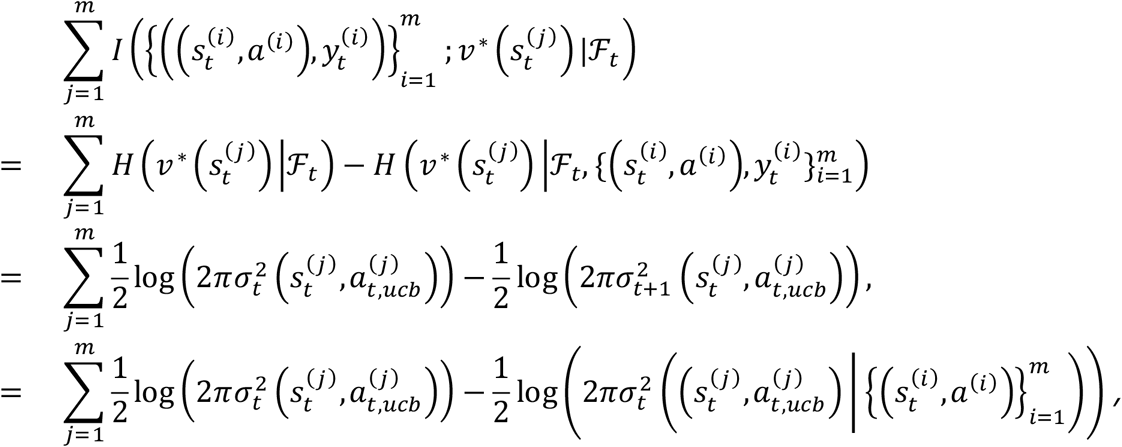

where the first equality follows from the property of mutual information (for any *X, Y, I*(*X; Y*) = *H*(*Y*) − *H*(*Y* ∣ *X*) where *H* denotes differential entropy, the second equality uses the definition of *v* ^*^ in (4), and the third equality uses the definition of 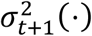, as well as the fact that the posterior standard deviation is independent of the measurement outcomes 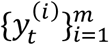 . Since 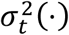 is independent of the choice of 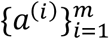, the objective in (3) is equivalent to

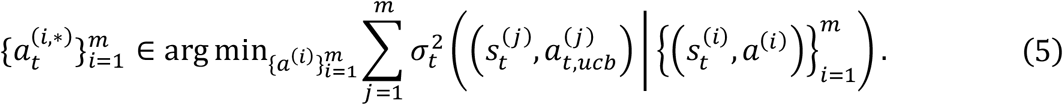

This can be interpreted as picking the actions 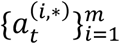 such that the uncertainty about the potential high-value points, i.e. 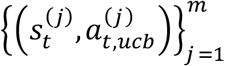, is reduced.

To further simplify (5), let 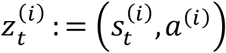, and denote 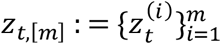. For any (*s, a*) pair, by (2), the following equality holds:

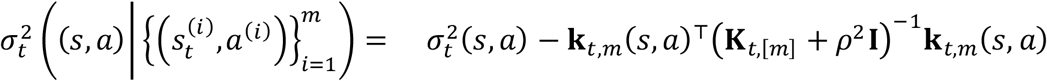

where

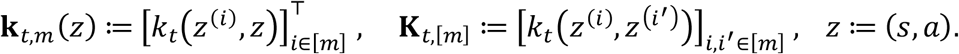

Hence, the objective in (5) can be further simplified to the equation in Line 10 of Algorithm 1:

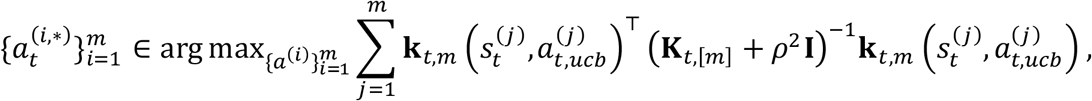

where

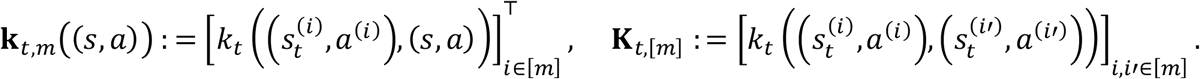

The actual implementation optimizes the objective in the Entropy Search section of Algorithm 1 by gradient descent.

In Lines 11 through 13 of Algorithm 1, the pseudotime change associated with the sampled actions at the next measurement day is measured and this data is added to the running dataset. Finally, when the experiment finishes, the optimal policy output by the algorithm is given in Line 15 of Algorithm 1.

## Data availability

All data that supporting the findings of this study are available within the paper and its Extended Data Figures.

## Code availability

Code for the parallel Bayesian Optimization algorithm of our paper is available at the following Github link: https://github.com/CoNG-harvard/bio_batch_BO.

## Acknowledgments

We acknowledge the support from NIH/NIDDK 1DP1DK130673 (J.L.); NIH/ NSF ECCS-2038603 (J.L., N.L., J.D., and R.Lee); NIH/NLM 5R01LM014465 (J.L., N.L., and J.D.); NSF AI institute 2112085 (N.L.); and ARO ECP W911NF2310315 (J.D.).

## Author contributions

J.L., R.Liu, N.L. and Z.R. conceived the idea and designed the research. R.Liu fabricated and characterized electronics. Z.R., X.Z. and N.L. developed the algorithm. R.Liu, Q.L. and W.W. performed cell differentiation. R.Liu performed cell cultures and device integrations, R.Liu and Z.L. performed the immunofluorescence staining experiments. R.Liu and Z.R. designed the tissue culture platform with different stimulation conditions. R.Liu, X.Z. and Z.R. performed electrophysiological recordings. R.Liu performed stimulation experiments. X.Z., Z.R. and R.Liu analyzed electrophysiological recording and stimulation data. R.Liu, Z.R., X.Z., N.L. and J.L. prepared figures and wrote the manuscript. All authors have reviewed the final version of the manuscript. J.L. and N.L. supervised the study.

## Figures and figure captions

**Extended Data Fig. 1.**
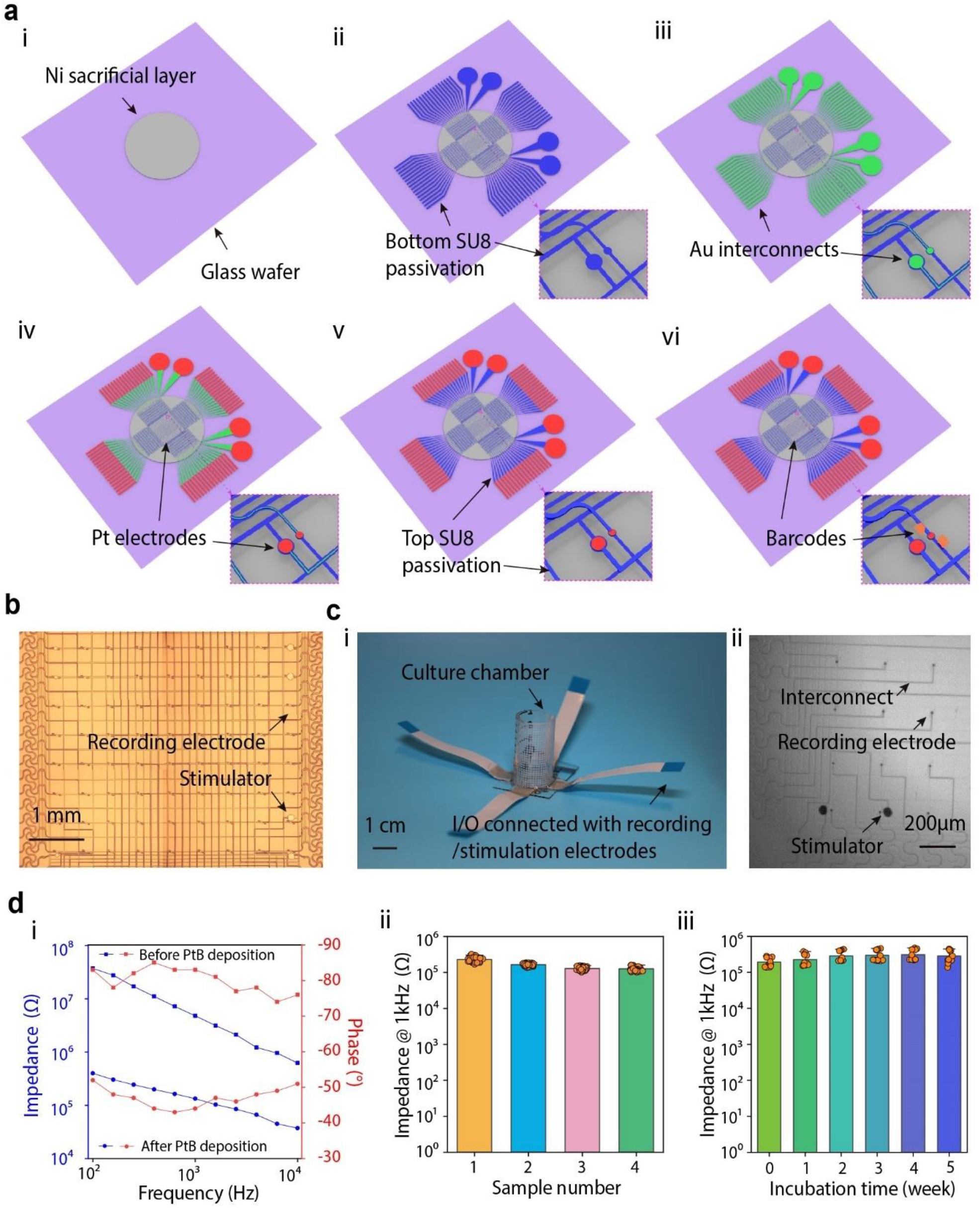
Fabrication of stretchable mesh electronics with sensing and stimulation electrodes. **a**, Schematic illustration of the fabrication process for stretchable mesh electronics. (i) Deposition of the Ni sacrificial layer on a glass wafer; (ii) Patterning of the bottom SU-8 passivation layer; (iii) Deposition and patterning of Cr/Au/Cr interconnects; (iv) Deposition and patterning of Pt electrodes; (v) Patterning of the top SU-8 passivation layer; (vi) Integration of electronic barcodes for electrode identification. **b**, Bright-field (BF) optical microscope image showing an overview of the 64-channel stretchable mesh electronics. **c**, (i) Photograph of the packaged stretchable mesh electronics with input/output (I/O) connections and culture chamber; (ii) Detailed view showing interconnects, recording electrodes and stimulators after releasing into saline water. **d**, Electrical characterization of stretchable mesh electronics. (i) Electrochemical impedance spectroscopy and phase response across frequencies for mesh electronics with and without Pt black; (ii) Statistics of impedance measurements at 1 kHz across different samples; (iii) Stability of impedance at 1 kHz over 50 days of incubation. Data are presented as means ± SEM.

**Extended Data Fig. 2.**
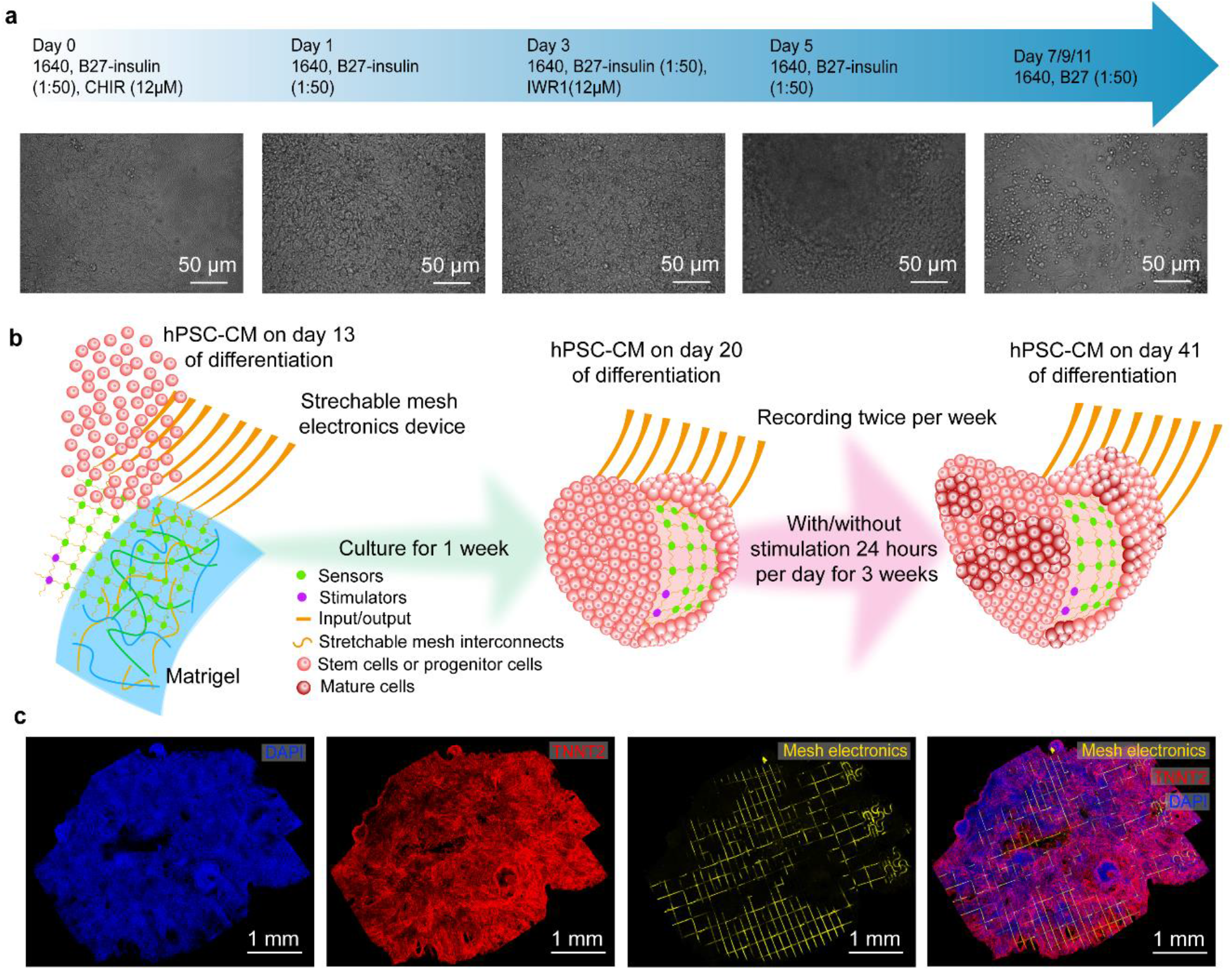
Integration of stretchable mesh electronics with hPSC-derived cardiac organoids. **a**, Timeline and phase contrast images of hPSC-CM differentiation. The differentiation protocol includes treatment with B27-insulin and CHIR at specified days. Day 0: Culture hPSC-CM in1640 medium with B27-insulin (1:50), and CHIR (12 µM); Day 1: Culture hPSC-CM in 1640 medium with B27-insulin (1:50); Day 3: Culture hPSC-CM in 1640 medium with B27-insulin (1:50), and IWR1 (12 µM); Day 5: Culture hPSC-CM in 1640 medium and B27-insulin (1:50); Days 7,9, and 11: Culture hPSC-CM in 1640 medium and B27-insulin (1:50). **b**, Schematics illustrating the integration of hPSC-CMs with stretchable mesh electronics. Left: Integration of hPSC-CMs with stretchable mesh electronics on day 13 of differentiation. Middle: Stimulation of hPSC-CMs begins on day 20 of differentiation; Right: Stimulation of hPSC-CMs concludes on day 41 of differentiation. **c**, 3D reconstructed fluorescent confocal microscopic images showing the structure and integration of mesh electronics with hPSC-CMs: Left: DAPI staining indicating cell nuclei. Middle Left: TNNT2 staining indicating cardiac troponin T in hPSC-CMs. Middle Right: Reflective mode imaging of mesh nanoelectronics, showing the mesh distribution. Right: Overlay of DAPI, TNNT2, and mesh, showing the successful integration and the spatial relationship between cells and electronics.

**Extended Data Fig. 3.**
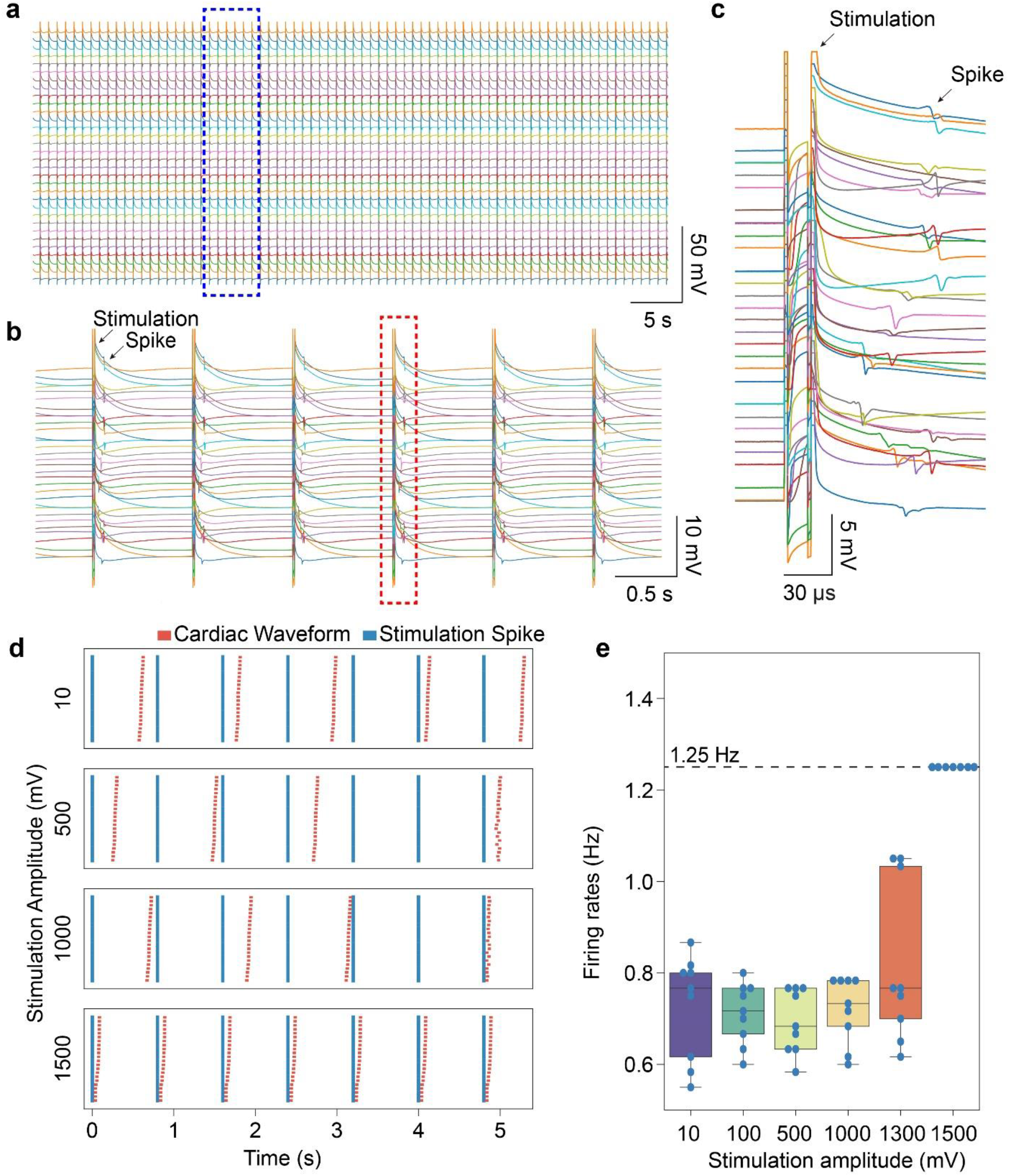
Stimulation and recording of human cyborg cardiac organoids. **a**, Representative raw traces of recording and stimulation. Each trace represents the electrical activity recorded from a specific electrode channel. **b**, Zoomed-in view of consecutive stimulation events, illustrating paced cardiac activity following the stimulation signals. **c**, Zoomed-in view showing detailed stimulation-induced cardiac spike characteristics for a single stimulation event across different channels. **d**, Raster plots showing cardiac activities under different stimulation amplitudes (10, 100, 500, 1000, and 1500 mV) over a 5-second period. Each plot corresponds to a different stimulation amplitude, illustrating the pacing pattern of electrical responses at 1500 mV of stimulation. **e**, Box plot of firing rates under different stimulation amplitudes. The firing rate fixed at 1.25 Hz under 1500 mV stimulation. Data points represent individual measurement. Box, 75% and 25% quantiles. Line, median. Whisker, 0% and 100% quantiles.

**Extended Data Fig. 4.**
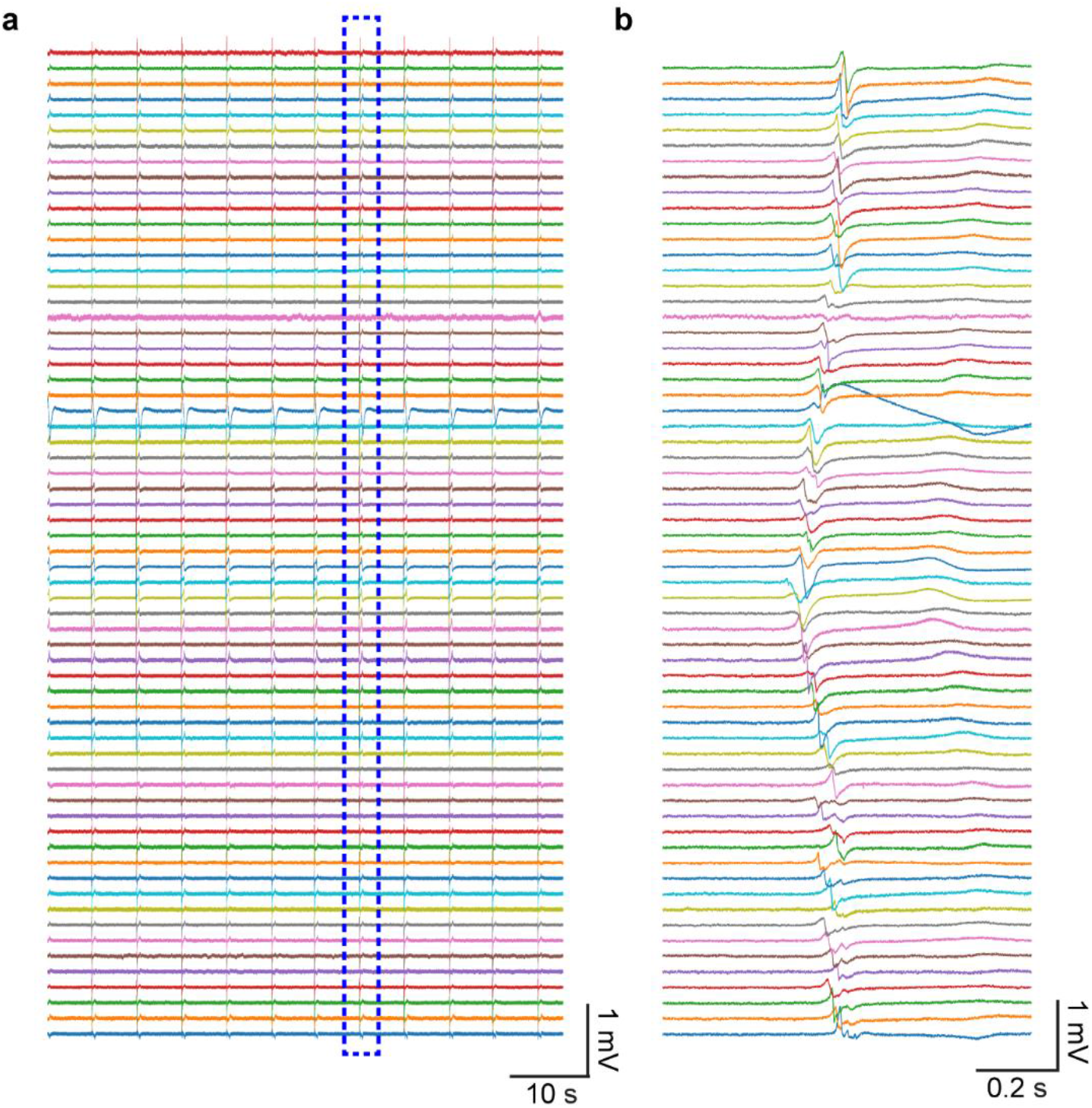
Representative electrical activity across a human cyborg cardiac organoid. **a**, Expanded view of continuous recording raw trace. **b**, Detailed view of cardiac activities recorded from multiple electrodes, illustrating the time delay of the cardiac activities at different locations.

**Extended Data Fig. 5.**
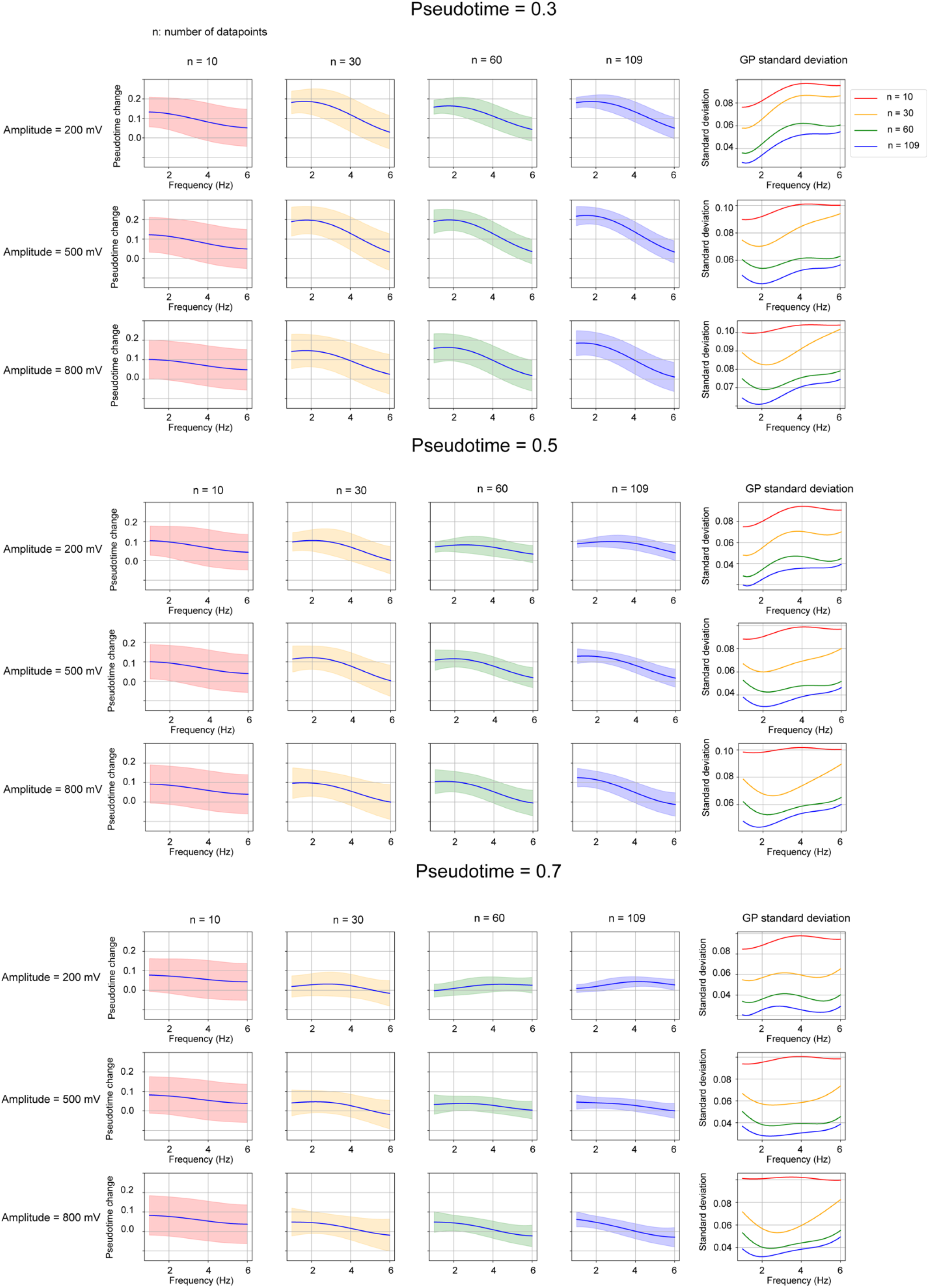
2D plots of Gaussian Process mean uncertainty for pseudotime change as the number of datapoints increases. The leftmost 4 panels of each row plot the change in predicted Gaussian Process mean and uncertainty (the shaded region corresponds to ± 1SD) for a fixed pseudotime (0.3, 0.5, 0.7) and amplitude (200 mV, 500 mV, 800 mV) as the frequency changes, with one panel for each of four number of datapoints (n = 10, 30, 60, 109). The rightmost panel of each row plots how the Gaussian Process standard deviation for pseudotime change varies with the frequency, with one curve for each number of datapoints.

**Extended Data Fig. 6.**
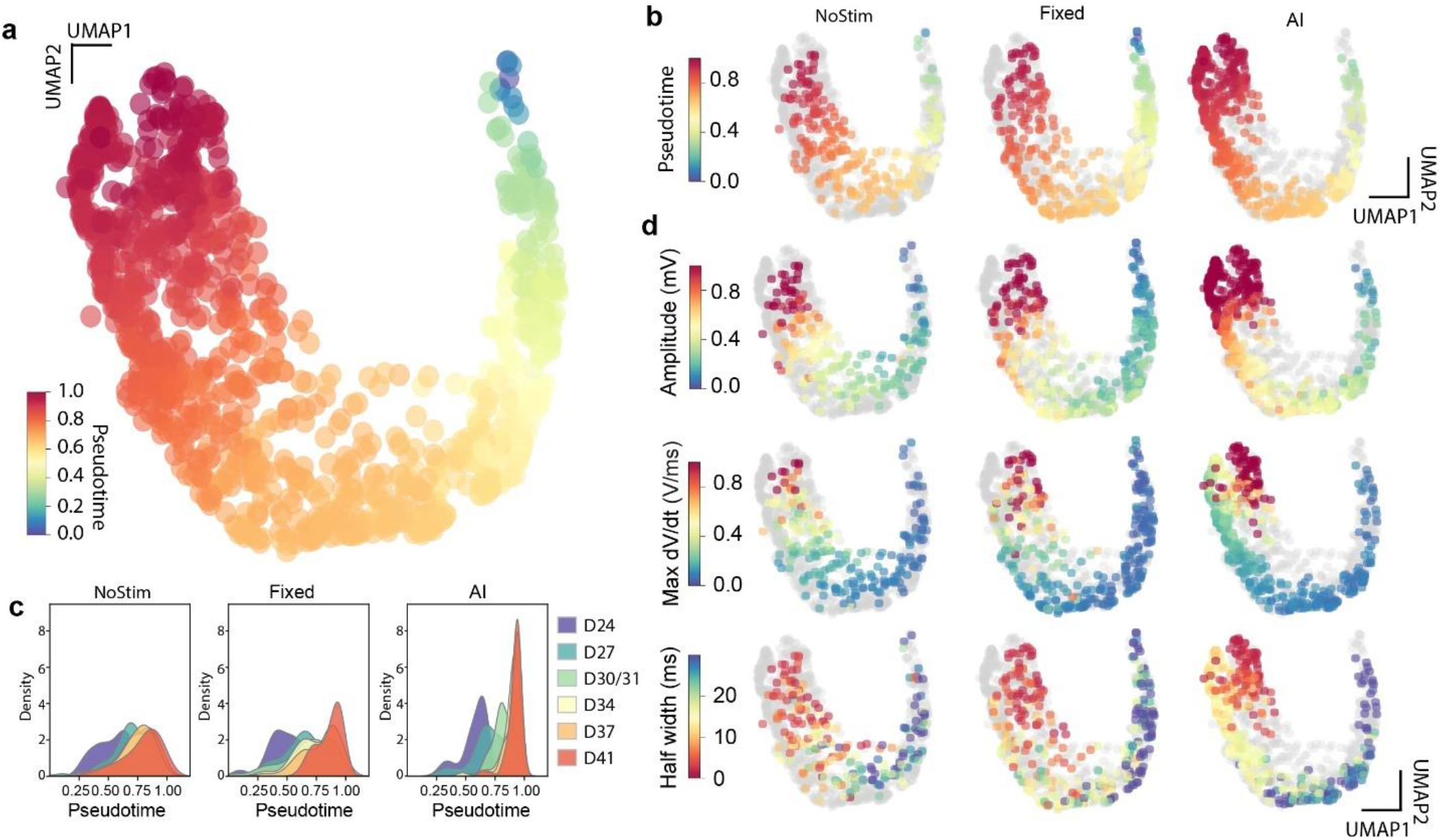
Comparative analysis of cell electrical activity in human cyborg cardiac organoids under different stimulation policies. **a**, Joint UMAP plots illustrating the distribution of pseudotime values for all six samples across two batches of experiments throughout the entire recording-stimulation period. Colors represent inferred pseudotime values. **b**, Joint UMAP for samples with three stimulation strategies: no stimulation (NoStim), fixed stimulation (Fixed), and AI-driven stimulation (AI). **c**, Density plots showing the distribution of pseudotime values across different differentiation days for the three stimulation conditions. Colors represent the differentiation days. **d**, Dot plots showing the distribution of electrical waveform characteristics, including amplitude, maximum dV/dt, and half-width, for each stimulation strategy and timepoints projected on the joint UMAP in (**a**). Colors represent the value of each characteristic. Each dot represents a channel-averaged electrical waveform.

**Extended Data Fig. 7.**
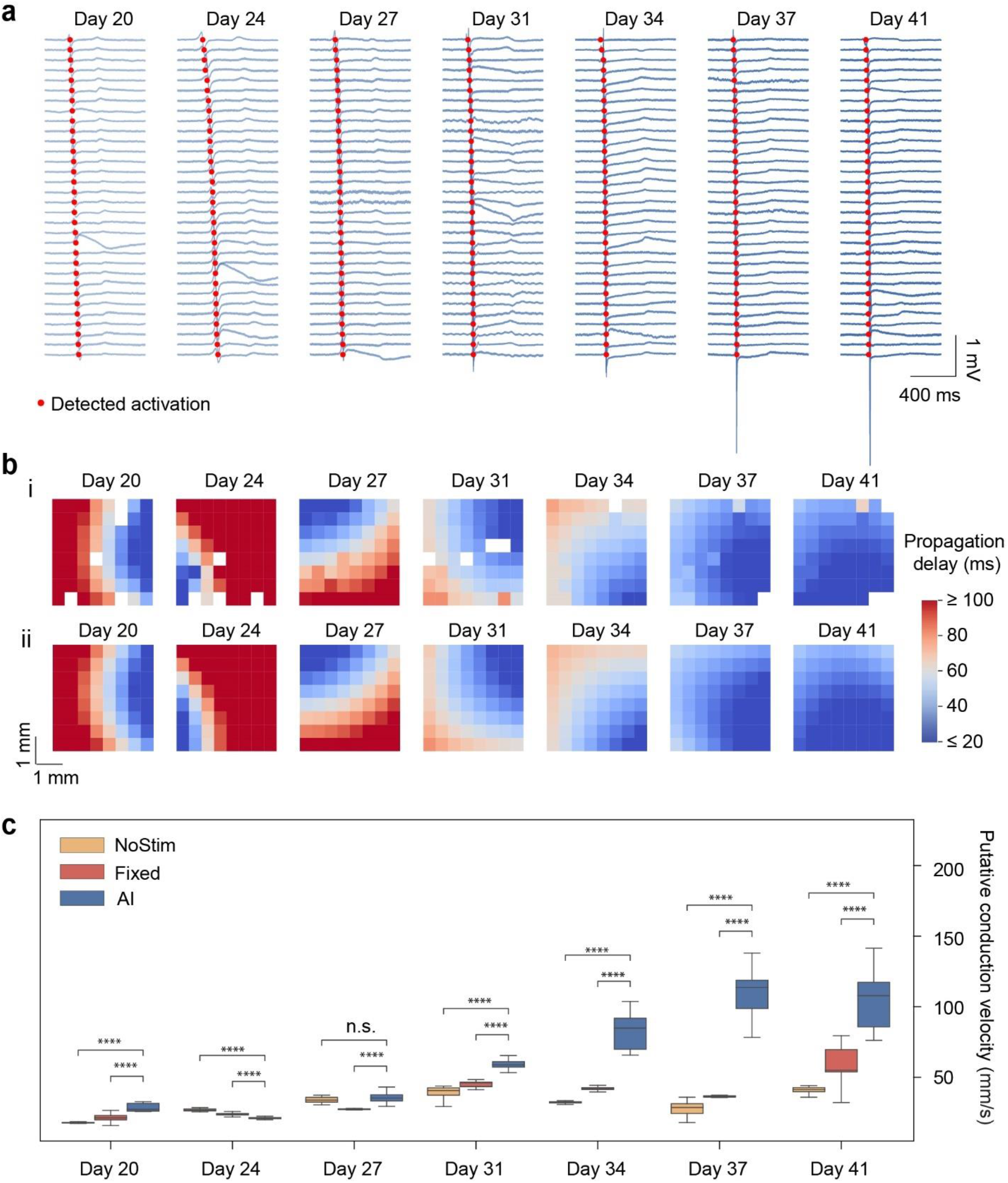
Comparative analysis of tissue-level electrical activity under different stimulation conditions. **a**, Representative raw voltage traces with activation time detection denoted by markers showing electrical activity in human cyborg cardiac organoids across various days of differentiation with AI-driven stimulation. **b**, Heatmaps illustrating the propagation delay of electrical signals across different channels of (i) raw map and (ii) modeled map of human cyborg cardiac organoids at various days of differentiation. The heatmaps use a color scale where red indicates longer delays and blue indicates shorter delays, showing the efficiency of signal propagation across the cardiac organoid tissue across various differentiation days. **c**, Box plots comparing the putative conduction velocity across different days of differentiation for control, fixed stimulation, and AI-driven stimulation groups. The plot shows significant differences in conduction velocity, with the AI-driven group displaying consistently higher values compared to no stimulation and fixed stimulation groups. **** p<0.0001, n.s. not significant, unpaired two tailed t-test. Box, 75% and 25% quantiles. Line, median. Whisker, the most extreme data point within the median ± 1.5 IQR.

**Extended Data Fig. 8.**
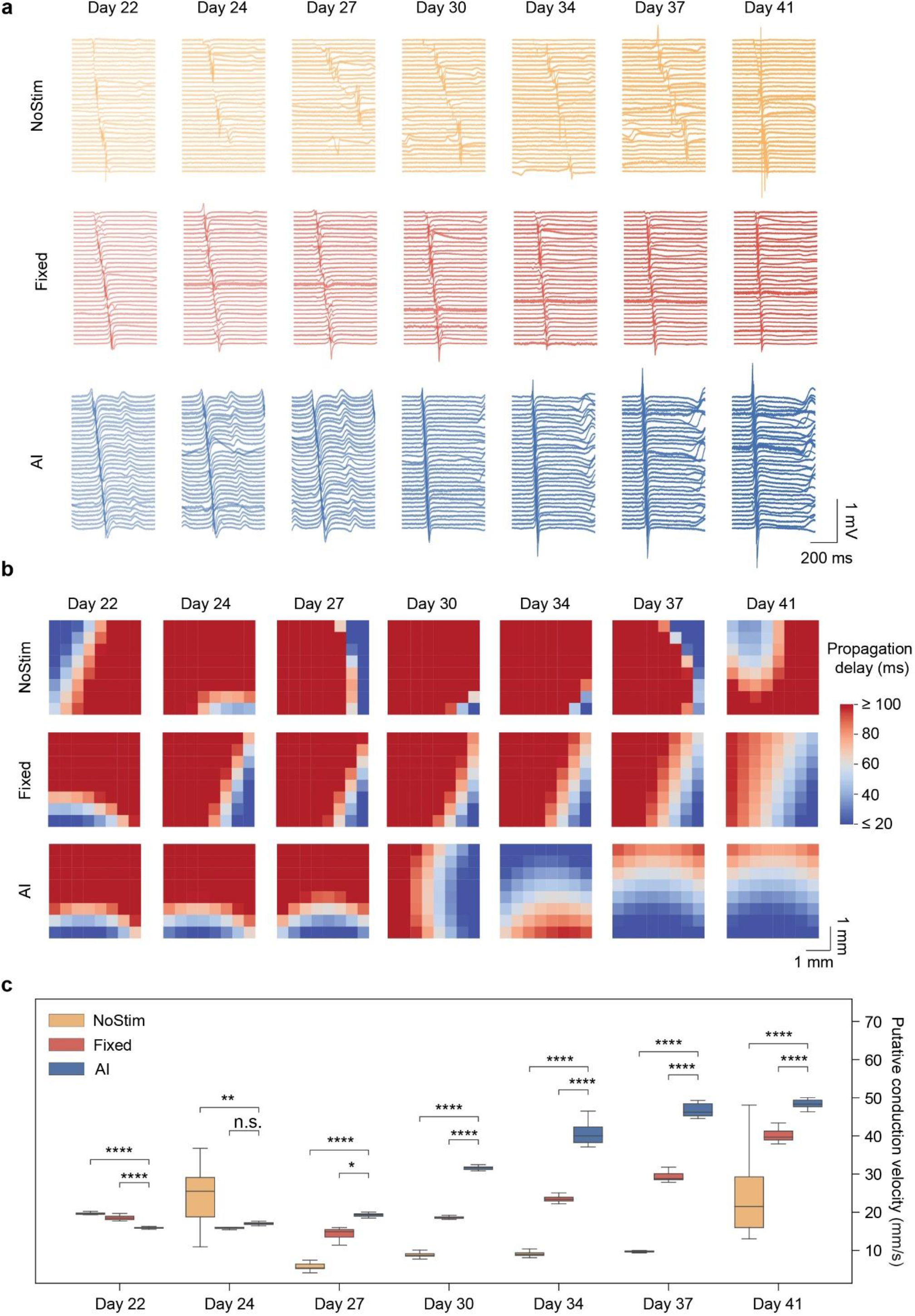
Biological replicates of tissue-level electrical activity under different stimulation conditions. **a**, Representative raw voltage traces showing the electrical activity of cardiac organoids under three different conditions across various days of differentiation. **b**, Heatmaps illustrating the propagation delay of electrical signals across different channels for no, fixed, and AI-driven stimulation groups at the same time points. Red indicates longer delays, while blue indicates shorter delays. **c**, Box plots comparing the propagation rate across different days of differentiation for control, fixed stimulation, and AI-driven stimulation groups. The plots show significant differences in conduction velocity, with the AI-driven group displaying consistently higher values compared to no stimulation and fixed stimulation groups. **** p<0.0001, ** p<0.01, * p<0.05, n.s. not significant, unpaired two tailed t-test. Box, 75% and 25% quantiles. Line, median. Whisker, the most extreme data point within the median ± 1.5 IQR.

**Extended Data Fig. 9.**
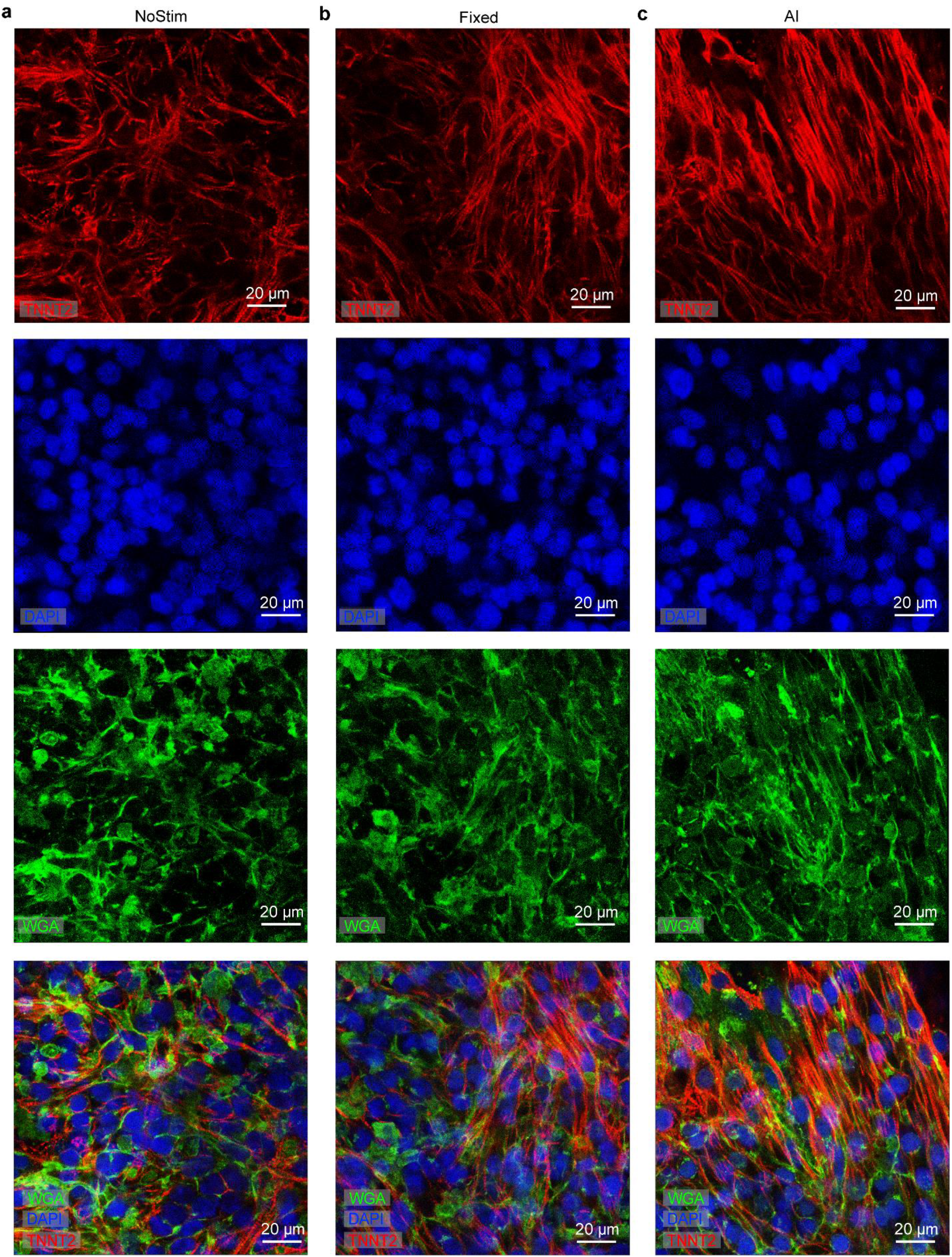
Immunofluorescence images of human cyborg cardiac organoids with no, fixed, and AI-driven stimulation policy. Immunofluorescence staining images of cardiac organoids with no stimulation (**a**), fixed stimulation policy (**b**), and AI-driven stimulation policy (**c**): Top row: Immunofluorescence staining for cardiac troponin T (TNNT2, red) indicating the sarcomere structure of cardiomyocytes. Second row: Immunofluorescence staining for DAPI (blue) highlighting cell nuclei. Third row: Immunofluorescence staining for wheat germ agglutinin (WGA, green) highlighting cell membranes. Bottom row: Immunofluorescence images of TNNT2 (red), wheat germ agglutinin (WGA, green), and DAPI (blue), showing sarcomere structures, cell membranes, and nuclei of cardiomyocytes, respectively.

**Table 1.** Summary of experimental groups and sample sizes for Figures 2, 4, 5, and Extended Data Figures 1, 3, 6, 7, 8. The table details the experimental conditions, including different development time points (days 20, 24, 27, 30/31, 34, 37, and 41), three stimulation strategies (NoStim, Fixed, AI), and their corresponding sample sizes (n values). Data are categorized by figure panels, with separate indications for specific experiments such as those involving inset analyses, batch effects, and different stimulation strategies. Each row indicates the number of samples (n) used under the specified conditions.

|  |  |  |  |  |  |  |
| --- | --- | --- | --- | --- | --- | --- |
| Figure 2 | d | e (Inset) | f/h/i/j<br>(Day 20) | f/h/i/j<br>(Day 24) | f/h/i/j<br>(Day 27) | f/h/i/j<br>(Day 31) |
| n value | 189 | 189 | 34 | 42 | 19 | 26 |
| Figure 2 | f/h/i/j<br>(Day 34) | f/h/i/j<br>(Day 37) | f/h/i/j<br>(Day 41) | k | l | m |
| n value | 19 | 20 | 29 | 189 | 189 | 189 |
| Figure 4 | d/e/f/h<br>(Day 24)<br>(Nostim) | d/e/f/h<br>(Day 27)<br>(Nostim) | d/e/f/h<br>(Day 30/31)<br>(Nostim) | d/e/f/h<br>(Day 34)<br>(Nostim) | d/e/f/h<br>(Day 37)<br>(Nostim) | d/e/f/h<br>(Day 41)<br>(Nostim) |
| n value | 43 | 42 | 42 | 39 | 29 | 38 |
| Figure 4 | d/e/f/h<br>(Day 24)<br>(Fixed) | d/e/f/h<br>(Day 27)<br>(Fixed) | d/e/f/h<br>(Day 30/31)<br>(Fixed) | d/e/f/h<br>(Day 34)<br>(Fixed) | d/e/f/h<br>(Day 37)<br>(Fixed) | d/e/f/h<br>(Day 41)<br>(Fixed) |
| n value | 92 | 58 | 66 | 40 | 37 | 40 |
| Figure 4 | d/e/f/h<br>(Day 24)<br>(AI) | d/e/f/h<br>(Day 27)<br>(AI) | d/e/f/h<br>(Day 30/31)<br>(AI) | d/e/f/h<br>(Day 34)<br>(AI) | d/e/f/h<br>(Day 37)<br>(AI) | d/e/f/h<br>(Day 41)<br>(AI) |
| n value | 89 | 88 | 86 | 93 | 76 | 79 |
| Figure 4 | g (Nostim) | g (Fixed) | g (AI) |  |  |  |
| n value | 233 | 333 | 511 |  |  |  |
| Figure 4 | i (Nostim)<br>Batch A | i (Nostim)<br>Batch B | i (Fixed)<br>Batch A | i (Fixed)<br>Batch B | i (AI)<br>Batch A | i (AI)<br>Batch B |
| n value | 10 | 28 | 11 | 29 | 47 | 32 |
| Figure 5 | c<br>(Day 20)<br>(Nostim) | c<br>(Day 24)<br>(Nostim) | c<br>(Day 27)<br>(Nostim) | c<br>(Day 31)<br>(Nostim) | c<br>(Day 34)<br>(Nostim) | c<br>(Day 37)<br>(Nostim) |
| n value | 144 | 18 | 23 | 38 | 31 | 19 |
| Figure 5 | c<br>(Day 20)<br>(Fixed) | c<br>(Day 24)<br>(Fixed) | c<br>(Day 27)<br>(Fixed) | c<br>(Day 31)<br>(Fixed) | c<br>(Day 34)<br>(Fixed) | c<br>(Day 37)<br>(Fixed) |
| n value | 121 | 176 | 23 | 34 | 39 | 29 |
| Figure 5 | c<br>(Day 20)<br>(AI) | c<br>(Day 24)<br>(AI) | c<br>(Day 27)<br>(AI) | c<br>(Day 31)<br>(AI) | c<br>(Day 34)<br>(AI) | c<br>(Day 37)<br>(AI) |
| n value | 17 | 11 | 18 | 28 | 41 | 112 |
| Figure 5 | c<br>(Day 41)<br>(Nostim) | c<br>(Day 41)<br>(Fixed) | c<br>(Day 41)<br>(AI) | f<br>(Nostim) | f<br>(Fixed) | f<br>(AI) |
| n value | 47 | 56 | 58 | 12 | 14 | 29 |
| Extended<br>Data<br>Figure 1 | d (ii)<br>(Sample 1) | d (ii)<br>(Sample 2) | d (ii)<br>(Sample 3) | d (ii)<br>(Sample 4) |  |  |
| n value | 62 | 62 | 63 | 62 |  |  |
| Extended<br>Data<br>Figure 1 | d (iii)<br>(Week 0) | d (iii)<br>(Week 1) | d (iii)<br>(Week 2) | d (iii)<br>(Week 3) | d (iii)<br>(Week 4) | d (iii)<br>(Week 5) |
| n value | 14 | 14 | 14 | 14 | 14 | 14 |
| Extended<br>Data<br>Figure 3 | e<br>Stimulation<br>amplitude<br>(10 mV) | e<br>Stimulation<br>amplitude<br>(100 mV) | e<br>Stimulation<br>amplitude<br>(500 mV) | e<br>Stimulation<br>amplitude<br>(1000 mV) | e<br>Stimulation<br>amplitude<br>(1300 mV) | e<br>Stimulation<br>amplitude<br>(1500 mV) |
| n value | 9 | 9 | 9 | 9 | 9 | 9 |
| Extended<br>Data<br>Figure 6 | a | b/d<br>(Nostim) | b/d<br>(Fixed) | b/d<br>(AI) |  |  |
| n value | 1077 | 233 | 333 | 511 |  |  |
| Extended<br>Data<br>Figure 7 | c<br>(Day 20)<br>(Nostim) | c<br>(Day 24)<br>(Nostim) | c<br>(Day 27)<br>(Nostim) | c<br>(Day 31)<br>(Nostim) | c<br>(Day 34)<br>(Nostim) | c<br>(Day 37)<br>(Nostim) |
| n value | 144 | 18 | 23 | 38 | 31 | 19 |
| Extended<br>Data<br>Figure 7 | c<br>(Day 20)<br>(Fixed) | c<br>(Day 24)<br>(Fixed) | c<br>(Day 27)<br>(Fixed) | c<br>(Day 31)<br>(Fixed) | c<br>(Day 34)<br>(Fixed) | c<br>(Day 37)<br>(Fixed) |
| n value | 121 | 176 | 23 | 34 | 39 | 29 |
| Extended<br>Data<br>Figure 7 | c<br>(Day 20)<br>(AI) | c<br>(Day 24)<br>(AI) | c<br>(Day 27)<br>(AI) | c<br>(Day 31)<br>(AI) | c<br>(Day 34)<br>(AI) | c<br>(Day 37)<br>(AI) |
| n value | 17 | 11 | 18 | 28 | 41 | 112 |
| Extended<br>Data<br>Figure 7 | c<br>(Day 41)<br>(Nostim) | c<br>(Day 41)<br>(Fixed) | c<br>(Day 41)<br>(AI) |  |  |  |
| n value | 47 | 56 | 58 |  |  |  |
| Extended<br>Data<br>Figure 8 | c<br>(Day 22)<br>(Nostim) | c<br>(Day 24)<br>(Nostim) | c<br>(Day 27)<br>(Nostim) | c<br>(Day 30)<br>(Nostim) | c<br>(Day 34)<br>(Nostim) | c<br>(Day 37)<br>(Nostim) |
| n value | 21 | 15 | 97 | 120 | 124 | 117 |
| Extended<br>Data<br>Figure 8 | c<br>(Day 22)<br>(Fixed) | c<br>(Day 24)<br>(Fixed) | c<br>(Day 27)<br>(Fixed) | c<br>(Day 30)<br>(Fixed) | c<br>(Day 34)<br>(Fixed) | c<br>(Day 37)<br>(Fixed) |
| n value | 19 | 17 | 18 | 19 | 20 | 25 |
| Extended<br>Data<br>Figure 8 | c<br>(Day 22)<br>(AI) | c<br>(Day 24)<br>(AI) | c<br>(Day 27)<br>(AI) | c<br>(Day 30)<br>(AI) | c<br>(Day 34)<br>(AI) | c<br>(Day 37)<br>(AI) |
| n value | 161 | 175 | 178 | 17 | 16 | 20 |
| Extended<br>Data<br>Figure 8 | c<br>(Day 41)<br>(Nostim) | c<br>(Day 41)<br>(Fixed) | c<br>(Day 41)<br>(AI) |  |  |  |
| n value | 39 | 49 | 22 |  |  |  |

## References

1 Lacour, S. P., Courtine, G. & Guck, J. Materials and technologies for soft implantable neuroprostheses. Nature Reviews Materials 1, 1–14 (2016).

2 Zhang, Y. et al. Printing, folding and assembly methods for forming 3D mesostructures in advanced materials. Nature Reviews Materials 2, 1–17 (2017).

3 Lee, G.-H. et al. Multifunctional materials for implantable and wearable photonic healthcare devices. Nature Reviews Materials 5, 149–165 (2020).

4 Zhao, C., Park, J., Root, S. E. & Bao, Z. Skin-inspired soft bioelectronic materials, devices and systems. Nature Reviews Bioengineering, 1–20 (2024).

5 Feiner, R. & Dvir, T. Tissue–electronics interfaces: from implantable devices to engineered tissues. Nature Reviews Materials 3, 1–16 (2017).

6 Jun, J. J. et al. Fully integrated silicon probes for high-density recording of neural activity. Nature 551, 232–236 (2017).

7 Steinmetz, N. A. et al. Neuropixels 2.0: A miniaturized high-density probe for stable, long-term brain recordings. Science 372, eabf4588 (2021).

8 Abbott, J. et al. A nanoelectrode array for obtaining intracellular recordings from thousands of connected neurons. Nature Biomedical Engineering 4, 232–241 (2020).

9 Abbott, J. et al. CMOS nanoelectrode array for all-electrical intracellular electrophysiological imaging. Nature Nanotechnology 12, 460–466 (2017).

10 Russell, S. J. & Norvig, P. Artificial Intelligence: a Modern Approach. (Pearson, 2016).

11 Nilsson, N. J. Principles of Artificial Intelligence. (Morgan Kaufmann, 2014).

12 Ertel, W. Introduction to Artificial Intelligence. (Springer, 2018).

13 Holzinger, A., Langs, G., Denk, H., Zatloukal, K. & Müller, H. Causability and explainability of artificial intelligence in medicine. Wiley Interdisciplinary Reviews: Data Mining and Knowledge Discovery 9, e1312 (2019).

14 Park, Y. et al. Three-dimensional, multifunctional neural interfaces for cortical spheroids and engineered assembloids. Science Advances 7, eabf9153 (2021).

15 Li, Q. et al. Multimodal charting of molecular and functional cell states via in situ electro-sequencing. Cell 186, 2002–2017 (2023).

16 Feiner, R. et al. Engineered hybrid cardiac patches with multifunctional electronics for online monitoring and regulation of tissue function. Nature Materials 15, 679–685 (2016).

17 Gao, H. et al. Graphene-integrated mesh electronics with converged multifunctionality for tracking multimodal excitation-contraction dynamics in cardiac microtissues. Nature Communications 15, 2321 (2024).

18 Jordan, M. I. & Mitchell, T. M. Machine learning: Trends, perspectives, and prospects. Science 349, 255–260 (2015).

19 LeCun, Y., Bengio, Y. & Hinton, G. Deep learning. Nature 521, 436–444 (2015).

20 Wainberg, M., Merico, D., Delong, A. & Frey, B. J. Deep learning in biomedicine. Nature Biotechnology 36, 829–838 (2018).

21 Segers, V. F. & Lee, R. T. Stem-cell therapy for cardiac disease. Nature 451, 937–942 (2008).

22 Garbern, J. C. & Lee, R. T. Cardiac stem cell therapy and the promise of heart regeneration. Cell Stem Cell 12, 689–698 (2013).

23 Noble, D. The rise of computational biology. Nature Reviews Molecular Cell Biology 3, 459–463 (2002).

24 Guo, Z. et al. Diffusion models in bioinformatics and computational biology. Nature Reviews Bioengineering 2, 136–154 (2024).

25 Sutton, R. S. Reinforcement learning: An introduction. A Bradford Book (2018).

26 Mnih, V. et al. Human-level control through deep reinforcement learning. Nature 518, 529–533 (2015).

27 Neftci, E. O. & Averbeck, B. B. Reinforcement learning in artificial and biological systems. Nature Machine Intelligence 1, 133–143 (2019).

28 Mahmud, M., Kaiser, M. S., Hussain, A. & Vassanelli, S. Applications of deep learning and reinforcement learning to biological data. IEEE Transactions on Neural Networks and Learning Systems 29, 2063–2079 (2018).

29 Angermueller, C. et al. Model-based reinforcement learning for biological sequence design. In International Conference on Learning Representations. (2020).

30 Treloar, N. J., Braniff, N., Ingalls, B. & Barnes, C. P. Deep reinforcement learning for optimal experimental design in biology. PLOS Computational Biology 18, e1010695 (2022).

31 Wang, Y. et al. Self-play reinforcement learning guides protein engineering. Nature Machine Intelligence 5, 845–860 (2023).

32 Wang, Z., Xu, Y., Wang, D., Yang, J. & Bao, Z. Hierarchical deep reinforcement learning reveals a modular mechanism of cell movement. Nature Machine Intelligence 4, 73–83 (2022).

33 Castrignanò, A., Bardini, R., Savino, A. & Di Carlo, S. A methodology combining reinforcement learning and simulation to optimize the in silico culture of epithelial sheets. Journal of Computational Science 76, 102226 (2024).

34 Salatino, J. W., Ludwig, K. A., Kozai, T. D. & Purcell, E. K. Glial responses to implanted electrodes in the brain. Nature Biomedical Engineering 1, 862–877 (2017).

35 Bendall, S. C. et al. Single-cell trajectory detection uncovers progression and regulatory coordination in human B cell development. Cell 157, 714–725 (2014).

36 Haghverdi, L., Büttner, M., Wolf, F. A., Buettner, F. & Theis, F. J. Diffusion pseudotime robustly reconstructs lineage branching. Nature Methods 13, 845–848 (2016).

37 Plass, M. et al. Cell type atlas and lineage tree of a whole complex animal by single-cell transcriptomics. Science 360, eaaq1723 (2018).

38 Lin, Z. et al. Tissue-embedded stretchable nanoelectronics reveal endothelial cell–mediated electrical maturation of human 3D cardiac microtissues. Science Advances 9, eade8513 (2023).

39 Fawzi, A. et al. Discovering faster matrix multiplication algorithms with reinforcement learning. Nature 610, 47–53 (2022).

40 Komorowski, M., Celi, L. A., Badawi, O., Gordon, A. C. & Faisal, A. A. The artificial intelligence clinician learns optimal treatment strategies for sepsis in intensive care. Nature Medicine 24, 1716–1720 (2018).

41 Andrychowicz, O. M. et al. Learning dexterous in-hand manipulation. The International Journal of Robotics Research 39, 3–20 (2020).

42 Bertsekas, D. Neuro-dynamic programming. Athena Scientific (1996).

43 Fatehullah, A., Tan, S. H. & Barker, N. Organoids as an in vitro model of human development and disease. Nature Cell Biology 18, 246–254 (2016).

44 Vlachogiannis, G. et al. Patient-derived organoids model treatment response of metastatic gastrointestinal cancers. Science 359, 920–926 (2018).

45 Shi, Y., Inoue, H., Wu, J. C. & Yamanaka, S. Induced pluripotent stem cell technology: a decade of progress. Nature Reviews Drug Discovery 16, 115–130 (2017).

46 Pagliuca, F. W. & Melton, D. A. How to make a functional β-cell. Development 140, 2472–2483 (2013).

47 Garreta, E. et al. Rethinking organoid technology through bioengineering. Nature Materials 20, 145–155 (2021).

48 Ronaldson-Bouchard, K. et al. Advanced maturation of human cardiac tissue grown from pluripotent stem cells. Nature 556, 239–243 (2018).

49 Nunes, S. S. et al. Biowire: a platform for maturation of human pluripotent stem cell–derived cardiomyocytes. Nature Methods 10, 781–787 (2013).

50 Ng, W. L., Chan, A., Ong, Y. S. & Chua, C. K. Deep learning for fabrication and maturation of 3D bioprinted tissues and organs. Virtual and Physical Prototyping 15, 340–358 (2020).

51 Garnett, R. Bayesian optimization. (Cambridge University Press, 2023).

52 Hennig, P. & Schuler, C. J. Entropy search for information-efficient global optimization. Preprint at ArXiv, 1112.1217 (2011).

53 Wang, Z. & Jegelka, S. Max-value entropy search for efficient Bayesian optimization. In International Conference on Machine Learning. 3627–3635 (PMLR, 2017).

54 Ma, H., Zhang, T., Wu, Y., Calmon, F. P. & Li, N. Gaussian max-value entropy search for multi-agent Bayesian optimization. In IEEE/RSJ International Conference on Intelligent Robots and Systems (IROS). 10028–10035 IEEE, 2023).

55 Li, Q. et al. Cyborg organoids: implantation of nanoelectronics via organogenesis for tissue-wide electrophysiology. Nano Letters 19, 5781–5789 (2019).

56 Le Floch, P. et al. Stretchable mesh nanoelectronics for 3D single-cell chronic electrophysiology from developing brain organoids. Advanced Materials 34, 2106829 (2022).

57 Setty, M. et al. Characterization of cell fate probabilities in single-cell data with Palantir. Nature Biotechnology 37, 451–460 (2019).

58 Silver, D. et al. Mastering the game of Go with deep neural networks and tree search. Nature 529, 484–489 (2016).

59 Kitano, H. Computational systems biology. Nature 420, 206–210 (2002).

60 Sidford, A., Wang, M., Wu, X., Yang, L. & Ye, Y. Near-optimal time and sample complexities for solving Markov decision processes with a generative model. Advances in Neural Information Processing Systems 31 (2018).

61 Williams, R. J. Simple statistical gradient-following algorithms for connectionist reinforcement learning. Machine Learning 8, 229–256 (1992).

62 Williams, C. K. & Rasmussen, C. E. Gaussian Processes for Machine Learning. Vol. 2 (MIT press Cambridge, MA, 2006).

63 Desautels, T., Krause, A. & Burdick, J. W. Parallelizing exploration-exploitation tradeoffs in Gaussian process bandit optimization. Journal of Machine Learning Research 15, 3873–3923 (2014).

64 András, V. et al. Cardiac transmembrane ion channels and action potentials: cellular physiology and arrhythmogenic behavior. Physiological Reviews (2021).

65 Bayly, P. V. et al. Estimation of conduction velocity vector fields from epicardial mapping data. IEEE Transactions on Biomedical Engineering 45, 563–571 (1998).

66 Zhu, H. et al. Two dimensional electrophysiological characterization of human pluripotent stem cell-derived cardiomyocyte system. Scientific Reports 7, 43210 (2017).

67 Sutcliffe, M. D. et al. High content analysis identifies unique morphological features of reprogrammed cardiomyocytes. Scientific Reports 8, 1258 (2018).

68 Bengio, Y., Courville, A. & Vincent, P. Representation learning: A review and new perspectives. IEEE Transactions on Pattern Analysis and Machine Intelligence 35, 1798–1828 (2013).

69 Zhang, T. et al. Making linear mdps practical via contrastive representation learning. In International Conference on Machine Learning. 26447–26466 (PMLR, 2022).

70 Ren, T. et al. Spectral decomposition representation for reinforcement learning. Preprint at ArXiv, 2208.09515 (2022).

71 Ren, T., Ren, Z., Li, N. & Dai, B. Stochastic nonlinear control via finite-dimensional spectral dynamic embedding. In 62nd IEEE Conference on Decision and Control (CDC). 795–800 (IEEE, 2023).

72 Kim, D.-H. et al. Stretchable and foldable silicon integrated circuits. Science 320, 507–511 (2008).

73 Liu, J. et al. Syringe-injectable electronics. Nature Nanotechnology 10, 629–636 (2015).

74 Le Floch, P. et al. Stretchable Mesh Nanoelectronics for 3D Single-Cell Chronic Electrophysiology from Developing Brain Organoids. Advanced Materials 34, e2106829–e2106829 (2022).

75 Lian, X. et al. Robust cardiomyocyte differentiation from human pluripotent stem cells via temporal modulation of canonical Wnt signaling. Proceedings of the National Academy of Sciences 109, E1848–E1857 (2012).

76 Lian, X. et al. Directed cardiomyocyte differentiation from human pluripotent stem cells by modulating Wnt/β-catenin signaling under fully defined conditions. Nature Protocols 8, 162–175 (2013).

77 Ji, Z. & Ji, H. TSCAN: Pseudo-time reconstruction and evaluation in single-cell RNA-seq analysis. Nucleic Acids Research 44, e117–e117 (2016).

78 Street, K. et al. Slingshot: cell lineage and pseudotime inference for single-cell transcriptomics. BMC Genomics 19, 1–16 (2018).

